# Antisense transcription can induce expression memory via stable promoter repression

**DOI:** 10.1101/2024.03.06.583761

**Authors:** Verena Mutzel, Till Schwämmle, Svearike Oeverdieck, Lucija Librenjak, Benedikt Boesen, Melissa Bothe, Rutger AF Gjaltema, Ilona Dunkel, Gemma Noviello, Edda G Schulz

## Abstract

The capacity of cells to retain a memory of previous signals enables them to adopt unique cell fates and adjust to their surrounding environment. The underlying gene expression memory can arise from mutual repression of two genes, forming a toggle switch. Such mutual repression may occur at antisense loci, where two convergently oriented genes repress each other in *cis*. Under which conditions antisense transcription can generate expression memory remains poorly understood. To address this question, we combine mathematical modeling, genomics and a synthetic biology approach. Through simulations we show that stable memory can emerge, if both genes in an antisense pair transcribe through the convergent promoter and induce a stable repressive chromatin state. Genome-wide analysis of nascent transcription further supports antisense-mediated promoter repression with promoter-overlapping antisense gene pairs exhibiting mutually exclusive expression. Through constructing a synthetic antisense locus in mouse embryonic stem cells (mESCs) we then show that such a locus architecture can indeed maintain a memory of a transient stimulus. Mutual repression and the capacity for memory formation are elevated, when mESCs differentiate, showing that epigenetic memory is a cell type-specific property. Our finding that stem cells adapt their ability to remember stimuli as they differentiate might help to elucidate how stemness is maintained.

## Introduction

Multicellular life requires the ability to memorize past signals. During embryonic development, cells must progressively commit to distinct cell lineages and stably remember their identity long after the commitment-inducing signals are gone. Also adaptation to environmental conditions frequently requires a memory of transient cues. The underlying gene expression memory can be established at the level of transcription factor networks in *trans* or at an individual allele in *cis*. *Trans*-memory uniformly affects all gene copies in a cell, whereas *cis*-memory enables individual alleles to retain distinct transcriptional states. Binary cell-fate decisions based on *trans*-memory are often achieved through cross-inhibition between two fate-determining transcription factors, a network motif called a toggle switch (Ferrell, 2002; Zhou & Huang, 2011). *Cis*-memory can be generated by self-reinforcing chromatin modifications, for example established through the Polycomb system (Dodd *et al*, 2007). In addition, many *cis*-memory genes are regulated by antisense transcription. The resulting mutual repression of two overlapping, convergently oriented genes in an antisense pair could form a *cis*-acting toggle switch and generate *cis*-memory.

Genomic imprinting is a prime example for *cis*-memory, where parental origin determines the allelic expression state. Antisense transcripts contribute to maintaining allele-specific gene repression at many imprinted loci (Kanduri, 2016). Also the *Xist* locus, which governs X-chromosome inactivation, is controlled by an antisense transcript in mice called Tsix (Lee & Lu, 1999). After Xist-driven inactivation of one randomly chosen X chromosome, Xist continues to be expressed from the inactive X throughout life, thus maintaining a memory of the initial random decision. Using mathematical modeling, we have previously shown that antisense transcription at the *Xist/Tsix* locus could help to lock in alternative expression states at the two alleles (Mutzel *et al*, 2019). More recently, antisense transcription has been described as a general hallmark of genes exhibiting random monoallelic expression (Kravitz *et al*, 2023), where it might have a functional role in stabilizing allelic expression states.

Given the pervasive transcription of the genome, >30% of coding genes exhibit antisense transcription (Katayama *et al*, 2005), though the term encompasses various phenomena. Many mammalian promoters show both divergent and convergent antisense transcription, originating slightly upstream and downstream of the transcription start site (TSS), respectively, producing short (<500bp) unstable transcripts (Mayer *et al*, 2015; Brown *et al*, 2018; Core *et al*, 2008; Preker *et al*, 2008). Contrarily, regulatory antisense RNAs at imprinted gene clusters and Xist are longer (>10kb), typically spliced transcripts with their own promoter (Pelechano & Steinmetz, 2013), enabling responsiveness to stimuli independently from the sense gene.

A variety of mechanisms have been described to underlie antisense-mediated gene repression (Pelechano & Steinmetz, 2013). Besides sequence-specific modes of regulation, repression can occur through transcriptional interference, where transcription itself mediates the repression. Collisions of two RNA polymerases transcribing in opposite directions can result in removal of one or both polymerases from the DNA (Noe Gonzalez *et al*, 2021). The frequency of collisions largely depends on the length of the overlapping region and the transcription rates, with their importance for regulation *in vivo* remaining unknown. More potent repression by contrast might occur when the antisense gene is transcribed through the sense promoter. The antisense polymerase can interfere with the formation of the pre-initiation complex, leading to so-called sitting-duck-interference, and can alter chromatin modifications at the promoter. Transcriptional elongation promotes the establishment of repressive marks, such as H3K36me3, which in turn recruits DNA methylation (Sims *et al*, 2004; Baubec *et al*, 2015; Teissandier & Bourc’his, 2017; Tufarelli *et al*, 2003; Pandey *et al*, 2008). This mechanism is thought to suppress transcription initiation within actively transcribed genes, but also has a functional role in antisense-mediated repression, e.g. at the *Xist* promoter (Ohhata *et al*, 2015). Chromatin-mediated repression can, in principle, function at lower transcription rates compared to sitting-duck-interference or polymerase collisions, since the repressed state can be maintained after the polymerase has passed.

Since antisense gene pairs can interfere with each other’s transcription through various transcription-mediated mechanisms, mutual repression could in principle establish a toggle switch at antisense loci. Expression of each gene, once established, would be self-reinforced through interference with its repressive antisense partner, thus stabilizing alternative expression states (Fig. 1A). It remains, however, largely unknown under which conditions antisense loci can maintain expression memory and whether it is facilitated by specific locus architectures or repression mechanisms.

**Figure 1.**
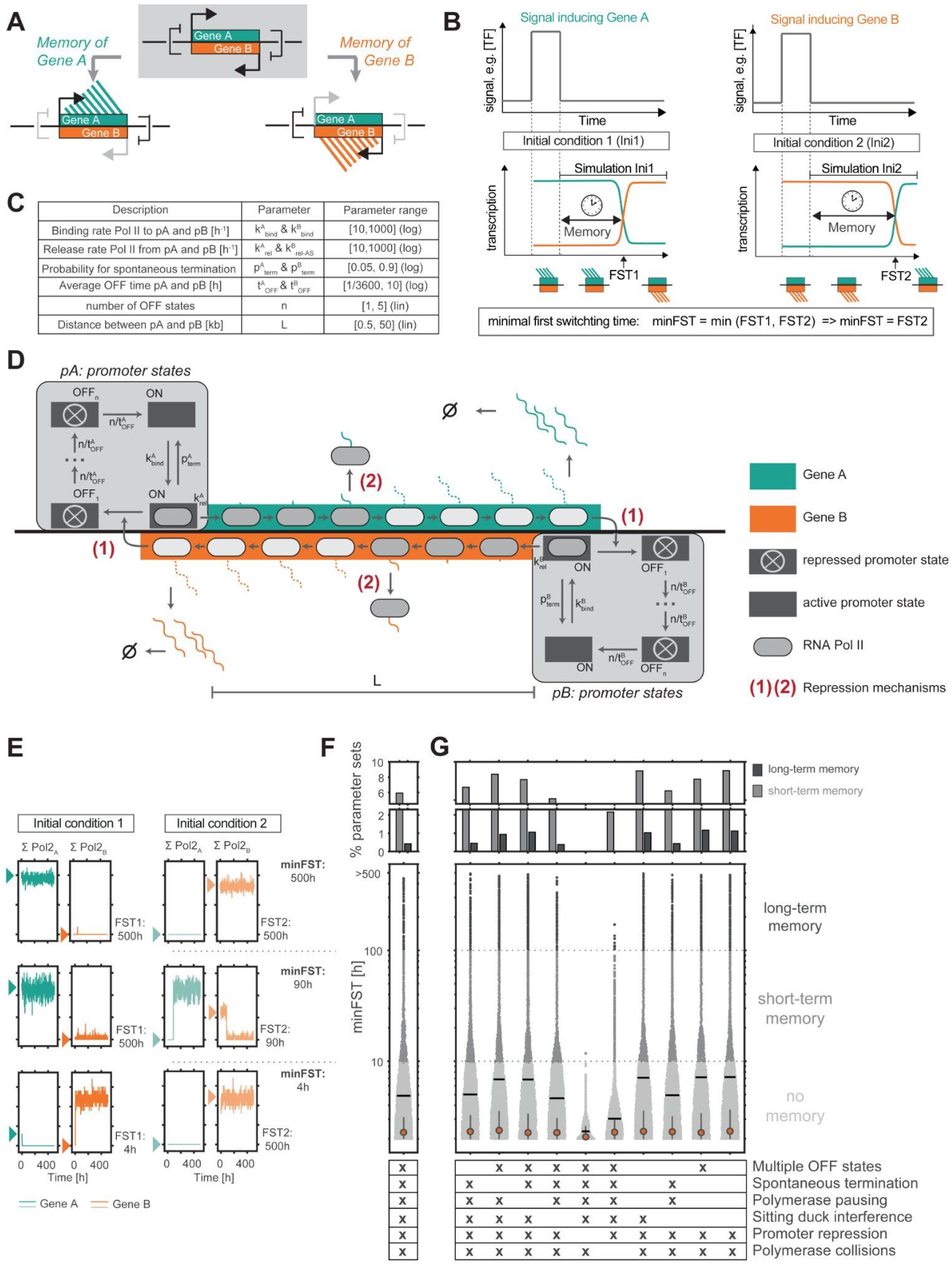
Mathematical model predicts that convergent promoters can memorize alternative expression states. A) Scheme of an antisense locus with two convergently oriented genes that mutually repress each other to form a toggle switch, that can maintain a memory of two alternative expression states. B) Schematic representation of allelic expression memory at an antisense locus. Left: In initial condition (Ini) 1 a transient transcription factor (TF) signal has induced Gene A leading to repression of gene B. Right: In Ini2 a transient signal has induced Gene B leading to repression of Gene A. The simulations E-G start once the transient signal is gone and measure the transcriptional memory, defined as the time that passes until the locus’ transcription state switches (FST1, FST2). minFST denotes the minimum of the two FSTs. C) Table summarizing the model parameters and tested ranges. D) Scheme of the mathematical model of antisense transcription. Pol II complexes can bind to both convergent promoters, when these are in the ON state, with distinct binding rates (k^A^, k^B^), and can then either be released into elongation (k^A^, k^B^), or spontaneously terminate transcription shortly downstream of the promoter (p^A^, p^B^). Elongating Pol II complexes move along the gene in convergent orientation. Mutual repression occurs through the act of transcription on the level of (1) promoter silencing by antisense transcription (ON-to-OFF transition induced by antisense Pol II at the promoter), and (2) transcriptional interference between two antisense Pol II complexes that occupy the same DNA segment, resulting in dislodgement of one Pol II from the DNA. E) Representative Gillespie simulation of transcription (sum of Pol II complexes on the gene) on strand A and B for example parameter sets with no (bottom), short-term (middle), and long-term (top) expression memory. Arrowheads indicate initial transcription state. Expression memory is quantified as minFST as shown in B. F) 35,000 parameter sets were classified into displaying no (light gray, minFST<10h), short-term (gray, 10h<minFST<100h), and long-term (dark gray, minFST>100h) expression memory for the model shown in C-D. Top: Percentage of parameter sets displaying short-term (gray) and long-term (dark gray) expression memory. Bottom: Distribution of minFST. G) Same as in F, but for a series of reduced models, where the process of transcription was simplified or mechanisms of repression between the two antisense strands were removed, as indicated. Each reduced model was simulated with 35,000 parameter sets. In F-G, small dots represent individual parameter sets, the red dots show the median, the horizontal black bars the mean and the whiskers the interquartile range.

Here, we combine mathematical modeling, genome-wide profiling and a synthetic-biology approach to elucidate the conditions under which antisense transcription can memorize a past signal. Through a series of simulations we show that antisense transcription could maintain expression memory up to several weeks, if both genes transcribe through each other’s promoter at a similar high rate and induce stable promoter repression. Using a time course data set of differentiating mouse embryonic stem cells (mESCs), measuring genome-wide nascent transcription we find that such antisense pairs with promoter overlap show indeed mutually exclusive expression, which becomes more pronounced during differentiation. We then build a synthetic antisense locus, capable of retaining short-term memory (∼1 day) in undifferentiated mESCs, which is stabilized during differentiation (> 4 days). Overall our interdisciplinary approach shows that antisense loci can memorize alternative expression states and that this property depends on the locus architecture, transcription rates and the global cellular state.

## Results

### Mathematical model predicts that expression memory can arise at antisense loci

We set out to investigate whether and to what extent mutual repression of two genes by convergent transcription can stably maintain alternative expression states (Fig. 1A+B). Specifically, we analyzed whether a transient signal (e.g. a transient change in transcription factor abundance or enhancer-promoter interaction frequency), inducing expression of one of the transcripts, could lead to a persistent change in the transcription state of the locus (*cis*-encoded memory).

We developed a mechanistic mathematical model of a generic antisense locus describing two convergent transcripts, Gene A and Gene B (Fig. 1C+D). The model accounts for transcription initiation (incl. pre-initiation complex assembly - k_bind_, release from promoter-proximal pausing - k_rel_ and pre-mature termination - p_term_), RNA polymerase II (Pol II) elongation, and RNA degradation of the antisense pair. Time-course experiments have suggested that activation of mammalian genes is a multi-step process (Harper *et al*, 2011; Suter *et al*, 2011; Coulon *et al*, 2013). To account for this observation, we modeled promoter transitions as an irreversible cycle of one transcriptionally active state (ON) and n inactive states (OFF) (Fig. 1D) (Zoller *et al*, 2015; Zhang *et al*, 2012).

The model assumes two mechanisms of repression between the convergent genes: (1) Transcription-dependent promoter repression through modification of the promoter-associated chromatin state and (2) transcriptional interference, occurring when Pol II complexes on two convergent strands meet. We differentiate between (i) encounters of two elongating Pol II complexes (collisions) and (ii) encounters between a promoter-bound Pol II and an elongating Pol II (sitting-duck-interference). Collisions result in dislodgement of one randomly chosen Pol II, while sitting-duck-interference always results in the removal of the promoter-bound Pol II. All of these mechanisms have been experimentally observed, and modeled in several previous studies (Sims *et al*, 2004; Baubec *et al*, 2015; Teissandier & Bourc’his, 2017; Brown *et al*, 2018; Gill *et al*, 2020; Murray *et al*, 2015; Nevers *et al*, 2018; Loos *et al*, 2015; Ward & Murray, 1979; Prescott & Proudfoot, 2002; Crampton *et al*, 2006; Hobson *et al*, 2012; Osato *et al*, 2007; Sneppen *et al*, 2005; Nakanishi *et al*, 2008; Mutzel *et al*, 2019).

To simulate a transient signal, which induces one of the genes (and thus indirectly represses the other), we start the simulation from two alternative initial expression states (Ini1, Ini2). In Ini1, Gene A is transcribed and Gene B is repressed, while in Ini2 a transient signal has induced Gene B leading to repression of Gene A (Fig. 1B). We performed the simulation in a stochastic manner using the Gillespie algorithm (only Pol II elongation was modeled deterministically) with more than 35,000 randomly chosen parameter sets. We combined a range of transcription strengths and locus features, to test whether – and under which conditions – the model could produce expression memory (Fig. 1C, Suppl. Table 1). The parameter ranges tested were based on previous experimental estimates (see methods section for details). For each parameter set, the simulation was performed 100 times.

The model stabilized alternative expression states up to several days for a subset of parameter values (example simulation in Fig. 1E, top). Notably, the transitions between the two expression states typically occurred in a switch-like manner, suggesting a bistable system. To quantify the stability of each expression state, we extracted the first switching times (FST), defined as the average time that elapses before the initially repressed gene becomes dominant (Gene B for Ini1 and Gene A for Ini2, Fig. 1B,E, see methods for details). To identify parameter sets that can stably maintain both states, we extracted the minimal FST (minFST) for each locus, given by the FST of the less stable state. We then classified parameter sets into those displaying no (minFST<10h), short-term (10h<minFST<100h), and long-term (minFST>100h) memory. While >90% of tested parameter sets exhibited no memory, 5.7% and 0.5% showed short-term or long-term memory, respectively (Fig. 1F).

To understand which inhibitory mechanisms and kinetic reactions were required to stabilize both states, we tested 10 model simplifications, removing one mechanism at a time (Fig. 1G). Removing antisense transcription-induced promoter repression completely abolished expression memory. Also, omission of transcriptional interference by collisions substantially reduced the average memory timescale and thus the percentage of parameter sets generating long-term expression memory. These results suggest promoter repression, and to a lesser extent polymerase collisions, as the main mechanisms underlying expression memory at antisense loci, which could stabilize alternative expression states several weeks under certain conditions.

### Antisense pairs with promoter overlap can stabilize alternative expression states for several days

We next wanted to understand the precise requirements for expression memory. Here, we focussed on the most simplified model, which retained promoter repression and polymerase collision, but contained only one promoter OFF state, did not account for pausing, premature termination and sitting-duck-interference (Fig. 1G, right, Fig. 2A). We resimulated the model with more parameter sets (119,000 sets in total), with additionally varying the frequency with which promoter repression and collisions occur (Fig. 2B). The strength of promoter repression was now controlled by two parameters: p_PR_ denotes the probability that a promoter switches to the OFF state, when transcribed by an antisense polymerase, and t_OFF_ defines the stability of the repressed state as the time that elapses before the promoter turns back ON. p_PR_ is assumed to be identical for both promoters, while t_OFF_ is a promoter-specific parameter.

**Figure 2.**
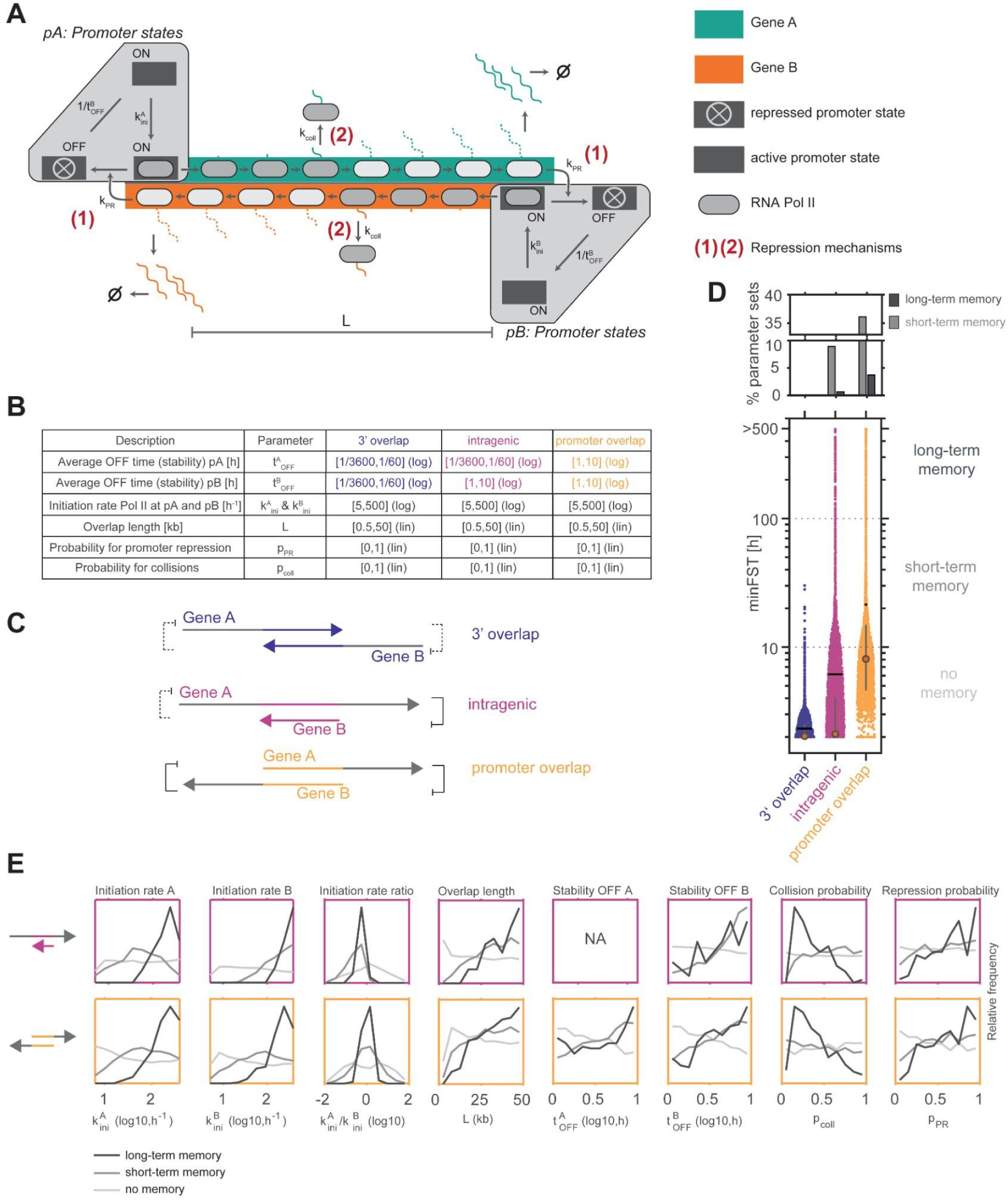
Long-term expression memory at antisense loci requires promoter repression. A) Scheme of maximally simplified model version with promoter repression (1) and polymerase collisions (2), one-step initiation, and a single promoter OFF state. B) Table summarizing parameter ranges tested in D-E and how antisense architectures are approximated by model parameters. C) Scheme depicting possible antisense transcription architectures including 3’ (blue), intragenic (pink) and promoter (yellow) overlaps. D) Distribution of minFST for different locus architectures (bottom) together with percentage of parameter sets displaying short-term (gray) and long-term (dark gray) expression memory (top). E) Distribution of parameter values for intragenic (top) and promoter overlap (bottom) architectures, across parameter sets displaying no (light gray), short-term (gray), and long-term (dark gray) expression memory.

We divided the simulated parameter sets into three categories that correspond to different locus architectures (Fig. 2B-C). Pairs, where the gene bodies overlap, but transcription does not proceed through the opposite promoter, were termed 3’ overlap. When transcription through the antisense promoter induces gene repression, the parameter set was categorized as ‘promoter overlap’, while pairs where one gene resides within another were termed intragenic. To compare the memory capacity between locus architectures, we again extracted the minFST for each parameter set. Pairs with promoter overlap had the highest memory potential, followed by intragenic overlaps, while 3’ overlapping pairs could not stabilize alternative expression states for extended time periods (Fig. 2D). These results are consistent with our previous finding that promoter repression is essential for long-term expression memory (Fig. 1G).

To better understand what allows certain parameter sets to stabilize alternative expression states, we analyzed the distribution of parameter values among the sets maintaining no, short-term, and long-term expression memory for the intragenic and promoter overlap architectures (Fig. 2E). The most stable sets exhibited frequent and stable promoter repression (high values for p_PR_ and t_OFF_), again pointing towards the importance of this repression mode. With respect to the second repression mechanism mediated by polymerase collisions, memory was most stable for parameter sets with a long overlap (high L, allowing more collisions), and a surprisingly low collision probability (p_coll_ ∼0.35). Although collisions must occur for stable memory, they should not be too frequent, potentially to still allow polymerases to reach the convergent promoter. Finally, stable memory was typically found for strong promoters (high k_ini_), probably required for efficient repression of the convergent gene through collisions and promoter repression. While for pairs with mutual promoter overlap, the initiation rates of the two promoters had to be tightly balanced for stable memory, at intragenic pairs the embedded gene (Gene B) required a stronger promoter, potentially because it had to rely solely on collisions to repress the convergent gene (Fig. 2E, top). Overall, the ability to repress the convergent transcript thus seems to be important for each strand to stabilize its transcribed state.

Taken together, our model analysis shows that mutual repression of antisense gene pairs could stably maintain transcriptional states for days to weeks, in particular if transcription-induced promoter repression is sufficiently strong. To test whether mutual repression at antisense loci indeed occurs *in vivo*, we next analyzed nascent transcription at antisense loci during the differentiation of mESCs.

### Genome-wide analysis detects skewed transcription at promoter-spanning antisense loci

To characterize endogenous antisense transcription in a genome-wide fashion, we re-analyzed a time-course experiment we had previously performed in female mESCs at days 0, 2 and 4 of differentiation by 2i/LIF withdrawal (Gjaltema *et al*, 2022). We reasoned that a joint analysis of different time points of this major cell state transition should allow us to observe dynamic changes in the expression states of antisense loci.

To detect antisense transcription we used Transient-Transcriptome sequencing (TT-seq) data, which profiles nascent transcription (Schwalb *et al*, 2016). TT-seq is based on short pulse-labeling of newly produced RNA combined with transcript fragmentation and therefore detects the entire transcribed region. Consequently, the method allows sensitive detection of lowly expressed and unstable transcripts, which are often involved in antisense transcription (Wery *et al*, 2018). To study antisense transcription beyond annotated genes, we first assembled a nascent transcriptome annotation for each time point from scratch (Fig. 3A, see methods for details), as done previously for such data (Shao *et al*, 2022). We segmented the genome into transcribed and non-transcribed 200 bp fragments for each strand. We detected putative TSSs as abrupt changes in nascent transcription (>10x fold change). As *bona fide* TSS, we then defined those that overlapped with a previously annotated TSS (GENCODE or mESC CAGE data) or with a promoter/enhancer state detected by chromatin segmentation (ChromHMM) based on measurements of DNA accessibility and active histone marks in the same cellular context (Frankish *et al*, 2019; Ernst & Kellis, 2012; Lizio *et al*, 2015, 2019; Gjaltema *et al*, 2022). Lastly, we removed any sporadic transcription without a verified TSS, trimmed the resulting transcription units (TUs) to a 10 bp resolution and merged those that overlapped on the same strand. In doing so, we recovered 17,028-18,481 TUs per time point, approximately half of which were not part of the commonly used GENCODE annotation (Fig. 3B, Suppl. Fig. 1A-D).

**Figure 3.**
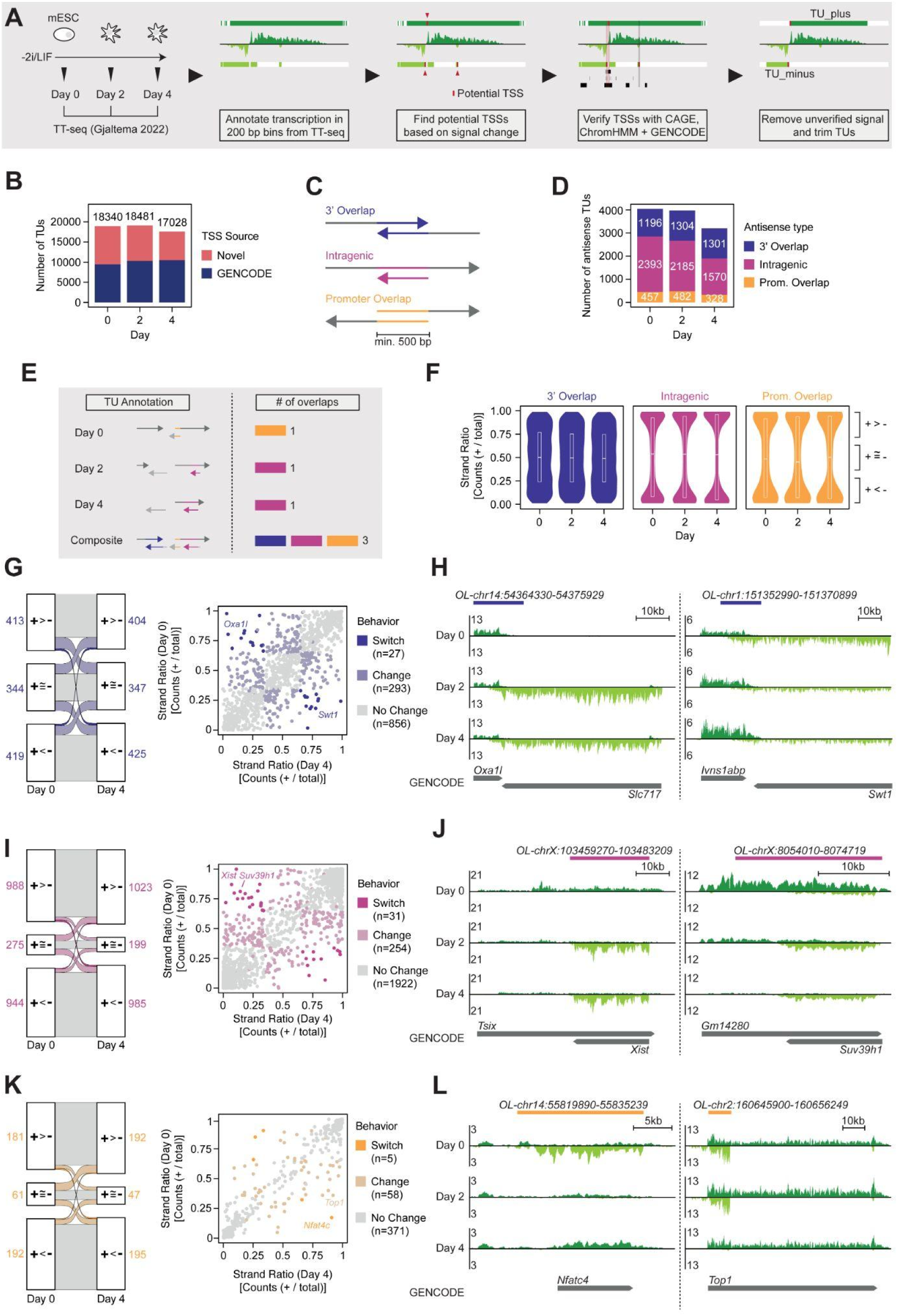
Genome-wide analysis of nascent transcription efficiently detects widespread antisense transcription. A) Scheme of nascent transcriptome assembly. TT-seq data from day 0, 2 and 4 of differentiation was used to annotate transcription genome-wide. Afterwards, potential TSSs were detected based on strong increases in transcription and verified by chromHMM, GENCODE and CAGE data (Frankish *et al*, 2019; Ernst & Kellis, 2012; Lizio *et al*, 2019, 2015). B) Number of annotated transcribed regions per time point. Transcribed regions whose TSS overlaps with a GENCODE transcript are shown in blue, others are shown in red. C) Scheme depicting possible architectures of antisense transcription including 3’ overlaps (purple), intragenic overlaps (pink) and promoter overlaps (yellow). D) Number of overlaps that were found using the assemblies of days 0, 2 or 4 of differentiation. The total number of overlaps per type is indicated on the respective bar in white. E) Scheme depicting the generation of a composite overlap set. The assemblies of 0, 2 and 4 days of differentiation were merged together and common overlaps called between all conditions. Overlaps shorter than 500 bp were discarded and intersecting overlaps combined. F) Strand ratio within the composite overlap set separated by day and overlap type. The ratio was calculated as the fraction between counts mapping to the plus strand. G) Sankey diagram and scatter plot depicting the strand ratio of 3’ overlaps at day 0 versus day 4. For each time point, an overlap was defined as biased, if transcription was significantly different between the two strands across n=2 biological replicates (Student’s T-test, Benjamini-Hochberg correction, FDR <= 0.1) and the strand ratio was >= 0.65 (plus bias) or <= 0.35 (minus bias). An overlap was annotated as ‘switch’, if it changed from one bias to the other and as ‘change’ if it changed from no bias to bias (or vice-versa). In the Sankey diagram, the total number of overlaps in each condition are depicted next to the boxes. H) Genome browser screenshot showing TT-seq data for two switching 3’ overlaps. Reads from the plus-strand are colored in dark green, while reads from the minus-strand are shown in light green. Overlap annotation is shown above the tracks. I) As in (G), but for intragenic overlaps. J) As in (H), but for intragenic overlaps. K) As in (G), but for promoter overlaps. L) As in (H), but for promoter overlaps.

To identify antisense loci, TUs on different strands that overlapped by at least 500 bp were detected and grouped into promoter, 3’ and intragenic overlaps (Fig. 3C-D). Intragenic and promoter overlaps were shorter on average (3542 / 2716 bp) compared to 3’ overlaps (8978 bp, Suppl. Fig. 1E). At each time point, 5,734-6,991 TUs were involved in at least one antisense pair. In mESCs and after 2 days of differentiation, we detected ∼4,000 antisense pairs, which was reduced to ∼3,200 at day 4. Interestingly, only the number of detected intragenic and promoter overlaps was reduced, while the number of 3’ overlaps remained stable (Fig. 3D). This means that loci where antisense transcription over a TSS is detected together with transcription from that TSS become less frequent upon mESC differentiation. One interpretation could be that antisense-mediated repression becomes more potent when mESCs differentiate.

Since our analysis so far treated each time point separately, we could not detect transcript pairs that are expressed in a mutually exclusive manner, where the dominant TU might switch upon differentiation. We therefore assembled a set of composite overlaps by combining the assemblies across the time course (Fig. 3E). After removing lowly expressed gene pairs, a set of 3817 antisense loci was identified, containing 434 promoter, 2207 intragenic and 1176 3’ overlaps. To investigate whether antisense transcripts show signs of mutual repression, we quantified to what extent they were expressed in a mutually exclusive manner. To this end, we defined a strand ratio as the fraction of TT-seq reads in the overlapping region that map to the plus strand (Fig. 3F). While transcription at intragenic and promoter overlap transcript pairs was strongly biased towards one of the strands (strand ratio close to 0 or 1), which became even more pronounced during differentiation, TUs in 3’ overlaps were often expressed at similar levels (strand ratio ∼0.5). A similar trend was observed when analyzing the time-point specific transcript annotation instead of the composite overlaps (Suppl. Fig. 1F). This finding suggests that transcription through a promoter indeed mediates gene repression, which seems to lead to switch-like behavior at antisense loci.

To test whether antisense-mediated repression might have a functional role during mESC differentiation, we next identified loci, where the dominant strand switched during the time course (Fig. 3G-L, Suppl. Fig. 1G-I, Suppl. Table 2). While the strand ratio remained unchanged for most antisense pairs, for each locus architecture we detected a small set of loci (5-31 antisense pairs) that switched the dominant strand (strand ratio >0.35 to <0.65 or vice versa). These include known antisense pairs such as *Xist/Tsix* and *Suv39h1/Suv39h1as* (Bernard *et al*, 2022; Schwämmle & Schulz, 2023), as well as loci that to our knowledge have not been implicated in antisense-mediated regulation in the past, for example *Nfatc4* and *Top1* (Fig. 3G-L). For these switching loci, antisense-mediated repression might be involved in their regulation during early mESC differentiation. Antisense pairs that are stable in our analysis might nevertheless switch in other cellular transitions.

### Promoter repression via antisense transcription correlates with increased DNA methylation

As our initial analyses identified mutually exclusive antisense transcription primarily at loci overlapping promoter regions, we examined the molecular mechanisms governing antisense transcription-mediated repression in more detail. We focused on 3’ overlaps to investigate polymerase collisions, and promoter overlaps to analyze antisense transcription-induced promoter repression. In the first step, we defined a set of overlap-free control TUs with matched length and nascent RNA expression for each antisense pair detected at day 0 for each locus architecture (Fig. 4A, Suppl. Fig. 2A, Suppl. Table 3) and compared their transcriptional activity throughout the time course to antisense TUs (Fig. 4B). TUs with 3’ overlap remained expressed at similar levels as the controls throughout the time course. TUs in promoter overlaps by contrast were more frequently downregulated upon differentiation compared to overlap-free controls (median fold change 0.30 compared to 0.95 in controls). When performing the same analysis for antisense pairs detected at day 4 with controls matched on day 4 expression, again controls exhibited little change across time points, while expression at antisense pairs increased over time (Suppl. Fig. 2B-C). The extent of upregulation, however, was smaller compared to the Day 0-centered analysis (median fold change 0.49 compared to 0.92). This observation suggests that antisense loci exhibit more dynamic expression than overlap-free genes. Moreover, the strong downregulation of antisense pairs expressed in undifferentiated cells might suggest that convergent transcription through a promoter becomes more repressive upon mESC differentiation. Antisense transcription in the gene body by contrast (3’ overlap), which cannot be mediated by promoter repression and might instead involve polymerase collisions, exhibited no sign of repressive activity in this analysis.

**Figure 4.**
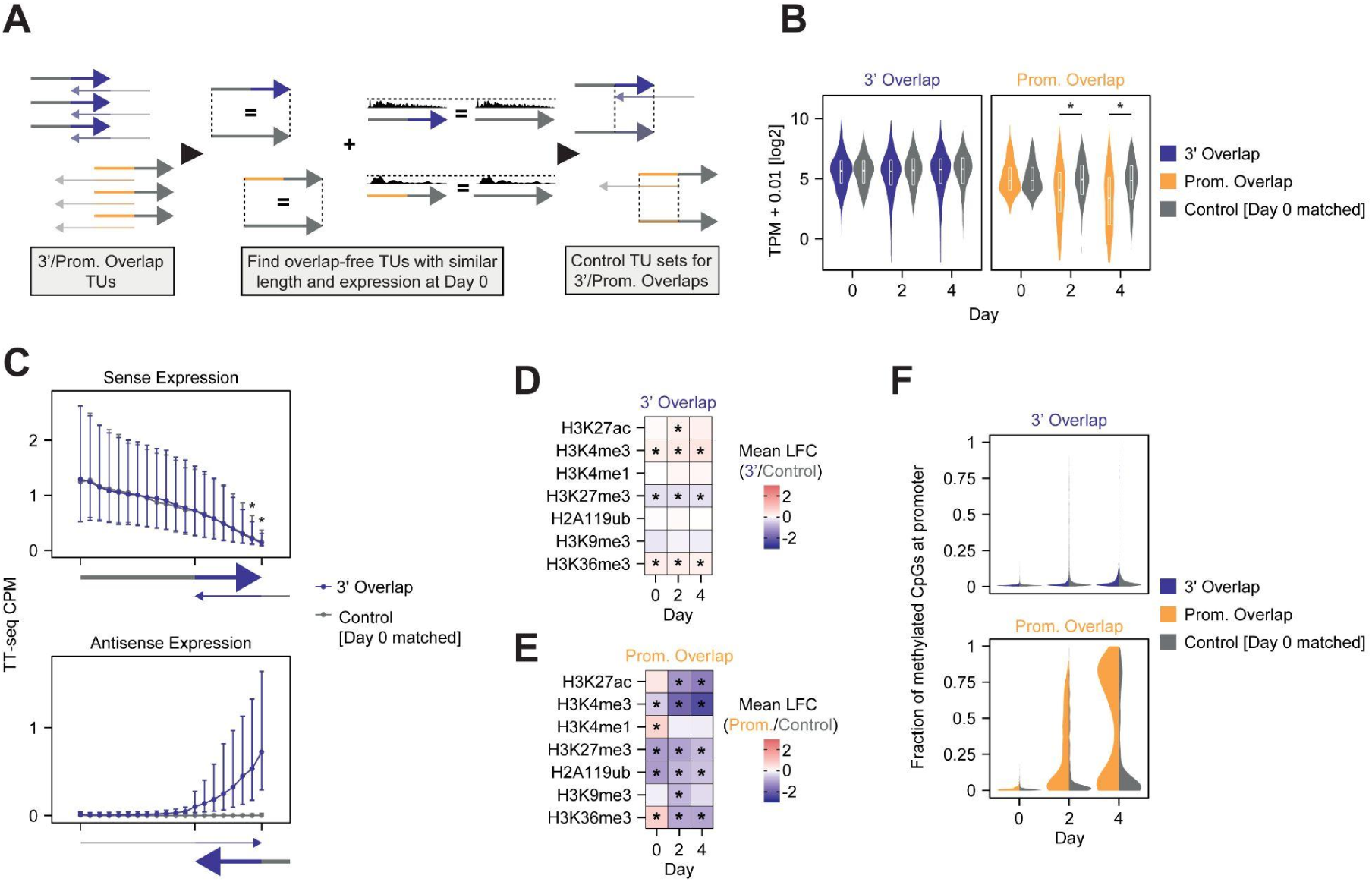
Promoter-overlapping antisense transcription primes for repression during differentiation. A) Scheme depicting the generation of a set of overlap-free control genes. B) Expression measured by TT-seq at 3’ and promoter-overlapping transcribed regions annotated based on transcription at day 0 in comparison to matched controls. Significance was assessed using a ranked Wilcoxon sum test (p<=0.01). C) Binned line plot showing sense or antisense expression of TT-seq data in 3’ overlap transcribed regions and mock-overlapping controls at day 0. Expression was quantified in 12 “free” bins and 8 “overlapping” bins. The big dots depict the median of all transcribed regions, while the upper and lower whiskers depict 3rd and 1st quartiles respectively. Significance was assessed using a ranked Wilcoxon sum test (p<=0.01). D-E) Heatmap depicting mean log2 fold change of different histone marks at promoters between 3’ overlapping transcribed regions (D) or promoter overlaps (E) compared to overlap-free controls. The CUT&Tag data was taken from (Gjaltema *et al*, 2022). * Significance was assessed using a ranked Wilcoxon sum test (p<=0.01). F) Violin plot depicting the percentage of promoter CpG methylation in 3’ or promoter overlapping TUs. Methylation levels for corresponding overlap-free controls (matched at day 0) are shown as gray diamonds. Promoters were defined as 500 bp upstream to 200 bp downstream of a TSS.

To more directly probe for transcriptional interference by polymerase collisions, we tested whether the TT-seq signal would decay in the overlapping part of 3’ overlap pairs more rapidly than control genes. We analyzed transcription activity along the transcripts in the overlapping and not overlapping region and compared them to overlap-free control TUs (for details see methods section). The distribution of transcription along the gene was very similar to control genes, albeit a slight reduction in the signal was observed towards the end of the transcript (Fig. 4C, Suppl. Fig. 2D). This suggests that transcriptional interference by polymerase collision is weak, or only present in a small subset of genes.

Having observed indications of antisense transcription-mediated promoter repression, in particular during differentiation, we asked what mechanisms could mediate the repression. We therefore examined the chromatin state at the promoters of antisense TUs. We re-analyzed a CUT&Tag dataset of histone marks that we had generated previously at the same time points as the TT-seq data (Gjaltema *et al*, 2022) and performed whole-genome bisulfite sequencing (WGBS) under the same conditions to analyze DNA methylation (see Suppl. Fig. 2E for example tracks).

As expected, we found only very small differences at promoters of 3’ overlapping TUs (where transcription does not extend through the promoter) compared to the overlap-free controls (Fig. 4D). For promoter-overlapping transcripts by contrast we found several differences (Fig. 4E-F). At day 0, promoters exhibited higher H3K36me3 levels, a mark deposited co-transcriptionally, which has previously been implicated in antisense-mediated repression (Fig. 4E) (Gerber & Shilatifard, 2003). At all time points, H3K4me3 levels were decreased in antisense promoters relative to controls (Fig. 4E). Among the repressive marks, only DNA methylation was enriched at antisense-transcribed promoters upon differentiation (Fig. 4F). H3K9me3 on the other hand, which is thought to become enriched at the *Xist* locus in response to antisense transcription (Ohhata *et al*, 2021; Gjaltema *et al*, 2022), was not globally associated with promoters repressed by antisense transcription, rather the opposite. The levels of H3K27me3 and H2AK119ub were even recorded below the overlap-free control group. As these marks are reportedly removed by transcription (Kaneko *et al*, 2014; Riising *et al*, 2014), this finding is not entirely surprising.

Taken together, our genome-wide analysis shows that antisense loci, where at least one partner transcribes through the promoter of the other one, exhibit signs of mutual repression and switch-like behavior. These findings are in line with our model analysis that proposed promoter repression to be the main determinant of memory, which is associated with switch-like behavior. With respect to polymerase collisions we find little experimental support in our genome-wide analysis, which could suggest that they occur rarely or only in a subset of loci. Finally, we find evidence for an increase in antisense-mediated repression strength during differentiation, potentially due to deposition of DNA methylation. As a consequence, mutual repression at antisense pairs becomes more pronounced during differentiation and we detect fewer pairs where both strands are transcribed simultaneously.

### A synthetic construct displays antisense transcription-mediated memory during mESC differentiation

Since our genome-wide analysis relies on correlations and cannot test for the existence of memory at antisense loci, we decided to use a synthetic biology approach to directly probe whether antisense transcription can maintain memory of a transient signal.

We designed a synthetic antisense locus that was stably integrated into the genome of female mESCs through a BxB1-dependent landing pad system, and named the resulting cell line TxSynAS (Fig. 5a). The synthetic nature of the construct allows us to monitor expression of both strands via fluorescent reporters and investigate the emerging regulatory responses independent of a functional role of the encoded transcripts or proteins. We initially tried to encode a reporter on each strand of the DNA sequence between two convergent promoters. We tested multiple different arrangements, with the antisense reporter encoded either in the 3’ UTR or 5’ UTR of the sense strand. We were, however, unable to find a design, where reporters from both strands could be detected by flow cytometry, maybe because the insertion of the antisense reporter sequence destabilized the transcripts. To circumvent this limitation, we encoded a sfGFP (hereafter called GFP) TU under the control of the EF1a promoter on the plus strand, and utilized a commonly used Doxycyline-inducible bidirectional promoter (pTRE-Tight) to control transcription in the antisense direction (Fig. 5A). In the antisense direction, Doxycycline (Dox) addition will drive convergent transcription across the GFP TU, and in the sense direction a fluorescent reporter (tdTomato) to monitor promoter activity. This strategy allowed us to bypass the need for the reporter to be encoded within the sense transcript. GFP and tdTomato were fused to a conditional FKBP destabilizing domain to accelerate protein turnover and thus temporal resolution of our measurements. However, the fluorescent signal of the destabilized reporters was too weak for reliable detection by flow cytometry. Therefore all experiments were performed in the presence of Shield1, which stabilizes the FKBP domain.

**Figure 5.**
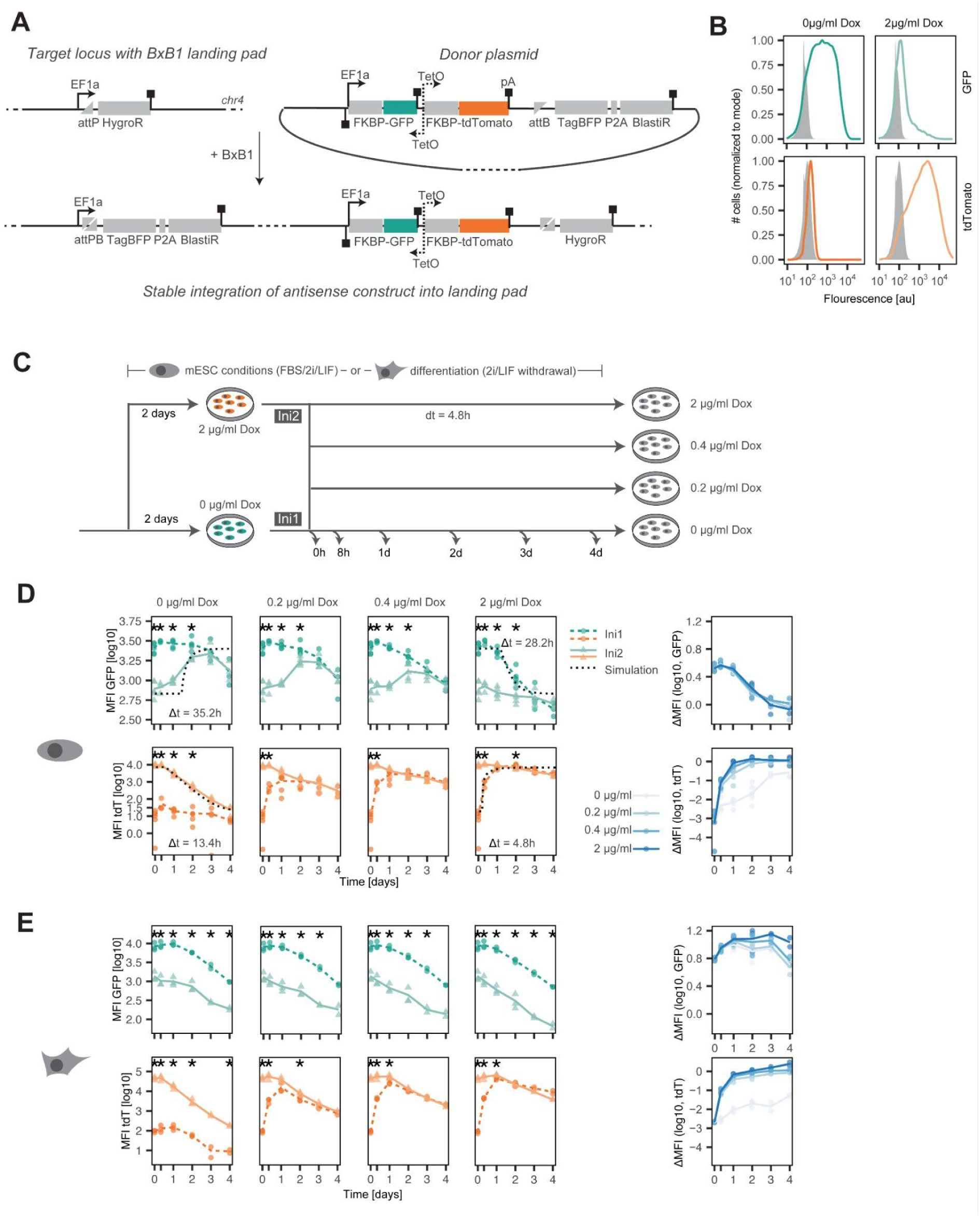
Antisense transcription can maintain expression memory in a synthetic antisense construct upon differentiation. A) Stable integration of synthetic antisense construct into genomic locus carrying a Bxb1-landing pad. Structure of the antisense construct developed in this study: human EF1a promoter drives transcription of a FKBP-coupled GFP. A bidirectional doxycycline-inducible TetO promoter drives convergent transcription across the GFP TU, and simultaneously transcribes a FKBP-coupled tdTomato reporter. Arrows and boxes indicate promoters and polyadenylation signals, respectively. B) Flow cytometry profiles of the TxSynAS line cultured for 2 days in the presence or absence of doxycycline. C) Scheme of doxycycline treatment profiles used in D and E. For two days, cells were either kept in medium without, or with high Dox concentrations to generate two different initial conditions Ini1 and Ini2. Cells were then cultured in four different Dox concentrations as indicated and reporter levels were assessed at the indicated time points by flow cytometry. D-E) Left: Background-corrected mean fluorescence intensity (MFI) of GFP (top) and tdTomato (bottom) in mESC medium FBS/2i/LIF (D) and upon differentiation by 2i/LIF withdrawal (E) for the experimental setup shown in C. Difference in expression levels between the two initial conditions was assessed with an unpaired Student’s t-test (p<0.05, asterisks). Black dotted lines represent the best fit of the ODE model. To quantify memory, we estimated a parameter Δt, indicated in the plots. Right: Time course of difference in reporter levels between the two initial conditions.

In the absence of Dox, cells express GFP, but not tdTomato (Fig. 5B, left). Treatment with Dox led to upregulation of tdTomato and a reduction in GFP expression (Fig. 5B, right), confirming that transcription from the convergent promoter represses expression of the sense strand, and that our assay can detect such antisense transcription-mediated repression.

Next, we set out to assess whether the locus would display expression memory, meaning that a transient increase or decrease in Dox concentration would induce a stable change in GFP expression. We treated cells for 2 days with the maximal amount of Dox (2µg/ml, Ini2) or without Dox (Ini1) and then cultured cells in 4 different Dox concentrations for another 4 days (Fig. 5C). We tested different concentrations since our model analysis suggested that memory might only arise for specific ratios of sense and antisense transcription rates. While the sense transcription rate is determined by the strength of the EF1a promoter, antisense promoter strength and thus also the promoter strength ratio, can be modulated by Dox. TdTomato levels rapidly converged towards the same expression level for both initial conditions (Fig. 5D bottom), which is expected since its bidirectional promoter is directly controlled by Dox. GFP levels in both populations also converged to the same level at the end of the experiment, albeit with slower dynamics (Fig. 5D, top).

To disentangle whether different dynamics of GFP compared to tdTomato indeed reflect expression memory or are driven by differences in protein stability, we used ordinary differential equation (ODE) models. We first estimated the rates of protein dilution through cell division and protein degradation for our GFP and tdTomato constructs, based on independent measurements and estimates from the literature (see methods section for details). We then tested whether model simulations could reproduce the data or whether an additional delay, potentially arising from expression memory, had to be assumed. We observed delays in the range of 30h for GFP, while the delays for tdTomato were substantially shorter (5-13h), and likely reflect incomplete Dox wash-out (Suppl. Table 4). The fact that the delay observed for GFP is >20h longer than the one observed for tdTomato, suggests that the synthetic antisense locus exhibits short-term expression memory in the range of one day in ESCs.

Since our genome-wide analysis had indicated that antisense-mediated promoter repression might be more potent upon differentiation, we assessed expression memory during differentiation. We repeated the memory assay with the antisense cell line upon withdrawal of 2i/LIF. We noted that even in the absence of Dox induction, GFP expression was gradually downregulated during differentiation (Fig. 5E, top, left, dark green). This might be due to downregulation of the EF1a promoter or silencing of the landing pad during differentiation, as the effect is conserved across different genomic integration sites (Suppl. Fig. 3). When comparing the effect of different initial conditions on the GFP reporter expression (-Dox and +Dox), we found that the sense reporter stably maintained alternative expression levels, determined by the initial condition, throughout the course of the experiment (4 days, Fig. 5E, top). This suggests that antisense transcription-induced repression of the sense reporter becomes stabilized upon differentiation. Conversely, tdTomato levels again rapidly converged towards the same level independent of the initial condition in most settings, confirming that transcription from the bidirectional promoter can still be potently induced (Fig. 5E, bottom). To normalize for the observed global downregulation of GFP expression upon differentiation, we quantified the difference in GFP levels between the two initial conditions over time (ΔMFI). In ESC conditions, ΔMFI converged towards zero, while it remained roughly constant upon differentiation (Fig. 5D-E, right), supporting stable memory induced by antisense transcription.

In summary, the experimental results support our theoretical model and genome-wide analysis, by pinpointing promoter repression as an essential mechanism for memory mediated by antisense transcription. Memory is stabilized upon mESCs differentiation, potentially due to enhanced DNA methylation, allowing maintenance of alternative expression states.

## Discussion

By combining three complementary approaches, we have dissected the conditions under which antisense transcription can give rise to expression memory. Through mathematical modeling and synthetic biology we showed that memory can be stable for several days, if RNA polymerases induce a long-lasting transcriptionally inactive chromatin state at the antisense gene promoter. This concept aligns with our observation of strong mutually exclusive activity at promoter-spanning convergent gene pairs from our genome-wide analysis, supporting the notion of transcription through the antisense promoter as repressive. Moreover, our findings suggest that the level of repression, and thus the potential for memory formation, is influenced by the cellular state. Repression appeared to be less pronounced in undifferentiated mESCs and strengthened as differentiation progressed, as evidenced by the detection of fewer co-transcribed antisense gene pairs with promoter overlap and stabilization of memory at the synthetic antisense locus.

Based on mathematical modeling we have established a quantitative framework to investigate the role of convergent transcription in generating *cis*-encoded expression memory. By systematically exploring various locus architectures and kinetic parameters, we discovered that enduring memory (lasting over 4 days) can be established when both genes exhibit high transcription rates (over 30 initiation events per hour) and trigger a repressed chromatin state with a lifetime in the range of hours. A recent genome-wide estimate of initiation rates in a mammalian cell line revealed a median rate of 30 h^-1^ and 9 h^-1^ for mRNAs and long non-coding RNAs, respectively (Gressel *et al*, 2019). In general, chromatin states can be stable for hours or even days (Lammers *et al*, 2020) and the antisense-transcription associated H3K36me3 mark has a lifespan for ∼1h in yeast (Lerner *et al*, 2020). Our analysis thus predicts antisense-mediated *cis*-memory to occur in a physiologically relevant regime.

Through construction of a synthetic antisense locus we showed that expression memory can arise from antisense transcription. The synthetic nature of our construct makes a regulatory role of the encoded transcripts highly unlikely, favoring a model in which the act of transcription mediates the establishment and maintenance of alternative expression states. While memory was only short-lived in mESCs, it became stabilized upon differentiation, in agreement with a previous study using a similar approach (Loos *et al*, 2015). The fact that antisense-mediated repression was potent in all conditions, but stable memory was restricted to differentiating cells, suggests that only a subset of repression mechanisms might be sufficiently stable to generate *cis*-memory. Work in yeast has established that antisense transcription induces histone deacetylation and loss of accessibility mediated by deposition of H3K36me3 (Brown *et al*, 2018; Gill *et al*, 2020; Nevers *et al*, 2018). Also, a study in mammalian cells, where Tsix, the antisense transcript of Xist was ectopically expressed in a cellular context where it is normally silent, revealed loss of active marks and accessibility as the earliest consequences of antisense transcription (Ohhata *et al*, 2021). These changes were, however, rapidly reversible, and only the more slow acquisition of repressive chromatin marks, such as H3K9me3 and DNA methylation, established a memory at Xist when Tsix transcription was turned off.

The fact that our synthetic locus maintained only a short-term memory in mESCs suggests that repression is short-lived in that cellular context. Our genome-wide analysis revealed that promoters transcribed in the antisense direction acquire H3K36me3, but not DNA methylation or H3K9me3. The basic antisense-mediated repression mode already present in yeast is thus already active in mESCs, but memory-forming mechanisms only become available upon differentiation. A similar observation was reported for another *cis*-memory system, mediated by Polycomb repressive complex 2 (PRC2). Transient PRC2 inhibition was fully reversible in mESCs, while a subset of target genes retained a memory of transient PRC2 loss, when the same experiment was performed in a more differentiated cell type (Holoch *et al*, 2021). What drives such acquisition of memory capacity remains largely unknown. An interesting correlation is that mESC differentiation is associated with a major increase in global DNA methylation levels (Kalkan *et al*, 2017; Schulz *et al*, 2024). Whether an increased activity of *de novo* DNA methyl-transferases indeed underlies *cis*-memory at antisense loci remains to be investigated.

An interesting avenue for future work will be to elucidate the specific roles of different *cis*-memory systems and how they interact with *trans*-memory. An intriguing hypothesis is that they operate at different time scales. At the *Xist* locus, for example, antisense transcription is only present transiently for a few days before other, potentially more stable, mechanisms, such as DNA methylation, ensure permanent repression (Sado *et al*, 2005; Shiura & Abe, 2019). Similarly, at the *FLC* locus in plants, antisense transcription can induce rapid silencing in response to external stimuli (temperature), which will later be consolidated by Polycomb repression (Nielsen *et al*, 2024). The role of antisense-mediated repression as compared to purely chromatin-based systems might thus be to establish *cis*-memory for a shorter time frame, while remaining responsive to external stimuli. How the coupling of different mechanisms ensures responsive, but robust memory remains to be investigated.

## Methods

### Simulations

#### Full antisense model

To simulate antisense transcription, initiation and elongation from two convergent promoters together with RNA degradation were combined into a mathematical model. Transcription initiation was modeled as a two-step process where Pol II complexes bind to the two convergently oriented promoters in a stochastic manner, and then are either released into productive elongation or spontaneously terminate transcription in the promoter-proximal region. Elongating Pol II complexes moved along the respective gene in a deterministic manner in steps of 100 nt length. All other reactions were simulated using the stochastic Gillespie algorithm. For the elongation and RNA degradation rates, experimental estimates were used, as explained in the next section. Transcription of a Pol II complex through the convergent promoter segment induced a switch to a transcriptionally inactive OFF state. To account for experimental data that suggest a peaked waiting time distribution in the silent state for mammalian promoters (Harper *et al*, 2011; Suter *et al*, 2011; Coulon *et al*, 2013), we modeled promoter reactivation as a multistep process, consisting of one active ON state where transcription can occur and n sequential inactive OFF states (Zoller *et al*, 2015; Zhang *et al*, 2012). When two convergent Pol II complexes occupied the same DNA segment, one randomly chosen Pol II was removed from the gene. If this segment was a promoter region, the promoter-bound Pol II was always removed from the DNA (sitting-duck-interference). Simulations were conducted in MATLAB_R2019b. The model was written in C++ and compiled into a MEX file that was called from the main MATLAB function. For parameter scanning, a compiled MATLAB script was executed in parallel on a computing cluster.

#### Simulating memory/maintenance of transcription state

To find parameter values that could maintain alternative transcription states at the simulated antisense locus, we first scanned a large parameter space. Degradation and elongation rates were set to fixed values based on previous experimental estimates of average rates in mammalian cells (Jonkers *et al*, 2014; Bartman *et al*, 2019; Zoller *et al*, 2015). All other parameters were randomly sampled within realistic parameter ranges (Supplementary Table 1, Sheet 1). We simulated >35,000 randomly sampled parameter sets.

To probe for conditions in which alternative transcription states at the antisense locus were stable, each allele was simulated twice starting from asymmetric initial conditions:

*Initial condition 1*: Gene A transcribed at steady state in the absence of antisense transcription-mediated repression, gene B not transcribed and promoter B OFF.

*Initial condition 2:* Gene B transcribed at steady state in the absence of antisense transcription-mediated repression, gene A not transcribed and promoter A OFF.

The steady state Pol II occupancy in the absence of transcriptional interference is calculated as:

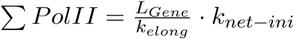

Where 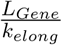 corresponds to the time that an elongating Pol II needs to transcribe the gene and 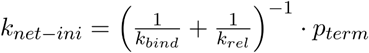 gives the net initiation rate, with 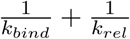 corresponding to the average time the promoter needs to produce one elongating Pol II. The steady state RNA level can then be inferred as the ratio of net production and degradation

rates: 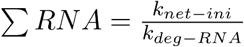

For each parameter set the antisense locus was simulated from both initial conditions for 500h in 100 realizations.

As a classifier for the stability of the initial transcription state the first-switching-time (FST) was extracted. The FST denotes the first time point at which the initially inactive gene dominated transcription within the overlap. It was defined as the first time point at which the ratio of the sum of Pol II complexes within the overlap, normalized to the relative promoter strength of the two genes, crossed 1:

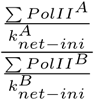

The FST was then averaged over all simulated alleles, and the minimum over the two initial conditions was used as a measure for transcriptional stability (minimal first-switching-time, minFST). A set was classified as exhibiting long-term memory if minFST > 100h, and as exhibiting short-term memory if 100h > minFST > 10h.

#### Model simplifications

To analyze which interference mechanisms and transcription reactions were strictly required for the generation of expression memory, we generated five reduced model versions, each either lacking one of the transcriptional interference mechanisms or simplifying transcription initiation or promoter reactivation into single-step reactions.

- In the model without promoter repression, passing Pol II complexes do not affect the state of the opposing promoter.
- In the model without collisions, sense and antisense Pol II complexes can bypass one another
- In the model without SDI, both Pol II complexes have the same probability to dislodge upon collision at the promoter segment
- To simplify promoter reactivation into a single-step reaction, we set the number of promoter OFF states N to 1.
- To simplify transcription initiation into a single-step reaction, we set k_rel_ = 10^5^ h^−1^ and p_term_ = 0 such that every Pol II that binds the promoter is immediately released into productive elongation. The new lumped one-step initiation rate k_ini_ was sampled between [5,500] h^−1^

Each simplified model was simulated with >35,000 randomly sampled parameter sets. The simplifications that did not substantially reduce the fraction of parameter sets displaying long-term transcriptional memory, were then hierarchically combined to obtain a maximally simplified model structure (see Fig. 2A and Suppl. Table 1, Sheet 2).

We resimulated the maximally simplified model with a larger number of parameter sets (119,000). Since it is unclear how frequently Pol II complexes collide in vivo, or induce a repressed promoter state, we included two additional parameters p_PR_ and p_Coll_ that modify the frequency of these events, respectively. Values for variable parameters were randomly drawn from uniform or logarithmic distributions (see Suppl. Table 1, Sheet 3).

#### Simulating different antisense locus architectures

Simulations were classified into the three different locus architectures based on the stability of the repressed promoter state. An average OFF period of <1 min was classified as unstable, while an average OFF period of >1 h was classified as stable. Those parameter sets with unstable repression of both promoters were categorized as 3’ overlaps, since promoter repression for both genes is essentially absent. Those sets with unstable repression of one, but stable repression of the other promoter were categorized as intragenic architecture, where only the promoter of the embedded gene can be repressed. Finally, all sets in which both promoters could be stably repressed were assigned to the promoter overlap category.

#### Quantification of memory in the synthetic antisense construct

To quantify the time scale of expression memory in the synthetic antisense construct, an ODE model was developed and model simulations were compared with the experimental data.

#### Quantification of reporter protein stability

As a first step, degradation rates of FKBP-GFP and FKBP-tdTomato in the construct were estimated. Wild type sfGFP and tdTomato protein half lives have previously been determined (Corish & Tyler-Smith, 1999; Kauffman *et al*, 2018). To estimate protein half lives of both reporters fused to the FKBP destabilization domain in the presence of 1µM stabilizing Shield1 ligand, we compared steady state expression levels of cell lines expressing the WT and FKBP-fused reporter version respectively, and found GFP to be destabilized 1.7x, and tdTomato 1.16x by fusion to the FKBP domain in presence of 1µM Shield1. With a half-life of 26h for the WT proteins this results in a half-life of 15.3h and 22.4h for FKBP-GFP and FKBP-tdTomato, respectively.

#### ODE model of reporter kinetics

An ODE model to describe GFP and tdTomato kinetics in the absence of expression memory was formulated assuming first-order degradation. The overall protein removal rate was assumed to depend on protein stability (degradation rate) and protein dilution due to cell division (dilution rate). The dilution rate is determined by the length of the cell cycle which in pluripotent mESCs has been measured to be roughly 12h (Waisman *et al*, 2019), while the degradation rate depends on the protein’s half-life (estimated above).

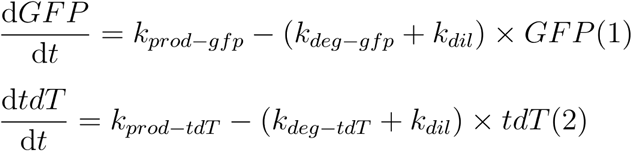

Since all measurements are relative, the production rates (k_prod-gfp_, k_prod_tdT_) were set arbitrarily to scale protein levels between 0 and 1. In the absence of Dox k_prod-gfp_ was set equal to the protein removal rate, and k_prod_tdT_ was set to 0. In the presence of maximal Dox levels k_prod-gfp_ was set to zero and k_prod_tdT_ was set equal to the tdTomato removal rate. for Equations (1) and (2) can then be solved analytically as follows

Dox removal:
1

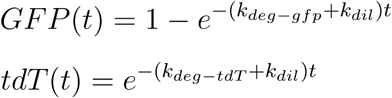

Dox addition:

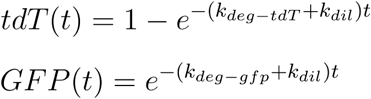

Analytical solutions of the ODEs were scaled to the experimentally determined time-averaged mean steady state expression level of cells cultured in the presence of no or 2µg/ml doxycycline in 2iL.

#### Quantification of memory at the synthetic antisense locus

To quantify whether and to what extent the synthetic antisense locus exhibits expression memory, we compared the kinetics predicted by the scaled ODE model to the flow cytometry expression data, assuming an immediate (Δt=0) or delayed (Δt>0) change in the production rate upon changing the Dox concentration in the culture media. For the offset Δt different values ranging from 0 to 48h were tested and the ODE solutions were compared to the experimentally measured GFP and tdTomato levels by calculating the sum of squared residuals (SRR) between the data and the model. Each experimental replicate was fitted separately to the ODE solution. The mean Δt values minimizing the SRR are reported in Figure 5. Suppl. Table 4 summarizes the mean and standard deviation of the fitted offsets.

### NGS data analysis

#### Nascent transcriptome assembly

To investigate antisense transcription genome-wide, we first generated a nascent transcriptome assembly based on TT-seq time course data set generated previously using the TXΔXic_B6_ mESC line (Gjaltema *et al*, 2022; Schwalb *et al*, 2016).

The data (GSE167356) was processed as described previously (Gjaltema *et al*, 2022). Additionally, blacklisted regions were removed from the BAM files using bedtools [intersect -v] (v2.29.2) (Quinlan & Hall, 2010). Subsequently, we merged the BAM files of individual replicates and used deeptools (v3.5.1) to count reads genome-wide in 200 bp bins on the plus and minus strands separately with options [multiBamSummary bins --genomeChunkSize 2407883318 --centerReads -bs 200] (Ramírez *et al*, 2016). Afterwards, the count tables were used as an input for the Genostan R package (2.24.0) to assign one out of 7 states to each bin depending on its transcription activity with options [initHMM(nStates = 7, method = “PoissonLogNormal”)] and [fitHMM(maxIters = 200)](Zacher *et al*, 2017). Per strand, the top 5 states were assigned as transcribed bins. To detect putative TSSs, we identified bins who showed a >10-fold TT-seq signal difference between the previous and the consecutive two bins. For an independent detection of potential TSSs we used previously generated ATACseq data, as well as CUT&Tag data of H3K27ac, H3K4me3 and H3K4me1 (GSE167350 and GSE167353) in the same cellular context to assign promoter and enhancer states using ChromHMM (v1.19) (Gjaltema *et al*, 2022; Ernst & Kellis, 2012). We then assigned each potential TSS detected in the TT-seq data as a true TSS only if the bin overlapped with a promoter/enhancer state from the ChromHMM analysis performed for the same time point, an annotated GENCODE TSS (Frankish *et al*, 2019) or a CAGE peak recovered by the FANTOM5 consortium (Lizio *et al*, 2015, 2019). The number of TSSs retrieved from either source is depicted in Suppl. Fig. 1A-C. To annotate the transcript associated with each verified TSS, the region was extended as far downstream as we could detect transcribed bins. Notably, we used the GENCODE annotation to fill holes between transcribed bins, if the TSS overlapped with an annotated GENCODE TSS. All other transcribed regions with an assigned TSS were removed from the analysis. To trim the annotated transcribed regions and increase the resolution, we counted reads in 10-bp bins covering the first and last 25% of each transcribed region using deeptools (v3.5.1) with options [multiBamSummary BED-file --centerReads] (Ramírez *et al*, 2016). Then we detected the first bin from the start of the transcribed region that surpassed 1/4 of the mean read counts in the region and designated it as the exact TSS. Lastly, we detected the first bin from the end of the TU that surpassed 1/2 of the mean read counts in the region and designated it as the corrected TTS. TUs that overlapped on the same strand (intragenic TSS) were merged. A complete list of all TUs for days 0, 2 and 4 of differentiation is provided in Suppl. Table 2.

#### Annotation of antisense overlaps

In order to find loci displaying antisense transcription, the generated (time-point-specific) annotations were used to find TU pairs on opposite strands that overlapped using GenomicRanges (v1.48.0) with options [findOverlaps ignore.strand = TRUE] (Lawrence *et al*, 2013). Subsequently, the overlaps were classified as promoter overlap, 3’ overlap or intragenic overlap depending on the architecture of the two TUs. Expression of the entire TUs and within the overlaps was quantified using Rsubread (v2.10.5) with options [featureCounts(allowMultiOverlap = TRUE)] (Liao *et al*, 2019). Data concerning the overlaps is supplied in Suppl. Table 2.

#### Annotation of composite overlaps

To create a composite set of overlaps throughout differentiation, we merged the assemblies of the three separate time points together. Afterwards, we once again annotated overlaps using GenomicRanges (v1.48.0) with options [findOverlaps(ignore.stand = TRUE)] (Lawrence *et al*, 2013). Subsequently, we removed all overlaps shorter than 500 bp and counted reads within each overlap at the three timepoints within the TT-seq data using Rsubread (v2.10.5) with [featureCounts(allowMultiOverlap = TRUE)] (Liao *et al*, 2019). We then calculated counts per million (CPM) for each condition and only kept overlaps with a minimum of three CPM at every time point on both strands combined. Lastly, we sorted the overlaps according to gene architecture as before and merged intersecting overlaps together.

#### Annotation of switching overlaps

In order to annotate overlaps which switched the dominant strand during differentiation, we first determined skewing in the composite overlap set at every time point and each of the 2 biological replicates, separately. To this end, we counted reads within the composite overlap set using Rsubread (v2.10.5) with [featureCounts(allowMultiOverlap = TRUE] and calculated CPM (Liao *et al*, 2019). For each time point, we used a two-sided Student’s T-test with Benjamini-Hochberg correction to detect antisense pairs where transcription was significantly different between the two strands (FDR <= 0.1). Among those, we defined an overlap as skewed if the ratio between the two strands [CPM_plus / (CPM_plus+CPM_minus)] was below 0.35 (minus-biased) or above 0.65 (plus-biased). Overlaps were annotated as “switch” if they changed the direction of the bias between the plus and minus strand throughout differentiation. Overlaps that were biased at one time point and balanced at the other were annotated as “change”. Overlaps that showed the same bias at all time points or stayed balanced throughout were annotated as “no change”. The results were visualized as Sankey diagrams and/or scatter plots using tidyverse (v1.3.2) and ggsankey (v0.0.99999) https://github.com/davidsjoberg/ggsankey (Wickham *et al*, 2019). A list of switch overlaps is given in Suppl. Table 2.

#### Generation of overlap controls

To study the effect of antisense transcription in 3’ or promoter overlaps in more detail we set out to generate an appropriate control set. To this end, we defined for each TU with a 3’ or promoter overlap an overlap-free region with similar length and expression. In detail, we filtered the day 0 or day 4 nascent transcriptome annotations for TUs longer than 2 kb with overlaps longer than 1 kb. Additionally, we removed every transcribed region with an overlap shorter than 5% or longer than 95% of the total transcript length. We then randomized the order of the overlap TUs and matched them to the free TUs in a loop, by matching on the maximal similarity in TPM and length. The free TU was then removed from the input, so that every TU could only be matched to an overlap TU once. We then created a “mock overlap” in each control matching the overlapped percentage of the partner TU. Lastly, the complete list of (mock) overlapping and free regions for promoter/3’ overlap transcription regions and controls were exported as BED files. Reads within the overlapping genes and the corresponding controls were quantified using Rsubread (v2.10.5) with [featureCounts(allowMultiOverlap = TRUE)] and expression throughout differentiation was quantified as TPM (Liao *et al*, 2019). Significance was assessed using a ranked Wilcoxon sum test (p<=0.05). A list of matched overlaps and control TUs are provided in Suppl. Table 3.

#### Generation of overlap lineplots

In order to investigate repression in 3’ overlaps, we compared expression along the 3’ overlap TUs and the corresponding, non-overlapping controls, to identify signs of premature termination due to polymerase collisions. To this end, we counted TT-seq reads using deepTools (v3.5.1) with [multiBamSummary scale-regions] in the overlapping and non-overlapping parts of the TUs separately (Ramírez *et al*, 2016). In the controls, which had a similar length (see previous section), a region of the same length as the overlap in the corresponding 3’ overlap TU was designated as a mock-overlap. Expression was quantified in 12 bins in the free part and 8 bins in the (mock-)overlap part. The number of bins was chosen according to the mean percentage of overlap length in the 3’ overlap control set (42.1%). Additionally, we also quantified expression in the antisense direction as a control. Significance was assessed using a ranked Wilcoxon sum test (p<=0.05).

#### Quantification of CUT&Tag data at promoters

The 3’ and promoter overlap sets detected at day 0, as well the corresponding control sets, were utilized to quantify H3K4me1, H3K4me3, H3K9me3, H3K27ac, H3K27me3, H3K36me3 and H2K119Aub data generated previously using CUT&Tag. The data (GSE167353) was processed as described previously (Gjaltema *et al*, 2022). Afterwards, counts were quantified from -500 bp to +200 bp around the TSSs using Rsubread (v2.10.5) with [featureCounts(allowMultiOverlap = TRUE)] (Liao *et al*, 2019). The data was normalized as CPM and visualized as a heatmap depicting the log2 fold change between the mean counts in the 3’/promoter overlap sets and the respective controls. Significance was assessed using a ranked Wilcoxon sum test (p<=0.01).

#### WGBS data processing

Adaptor as well as quality trimming of fastq files was performed using Trim Galore (v0.6.4) (https://github.com/FelixKrueger/TrimGalore (Martin, 2011) with the options [--clip_R1 10 --three_prime_clip_R1 5 --clip_R2 15 --three_prime_clip_R2 5 --paired]. Subsequently, reads were aligned to the mouse genome (mm10) with BSMAPz [-q 20 -u -w 100] (Xi & Li, 2009). The resulting bam files were sorted and low quality mapped reads were removed using samtools [view -q 10] (v1.10) (Li 2009). Next, duplicate reads were removed with Picard (v2.7.1) using the options [MarkDuplicates REMOVE_DUPLICATES=TRUE] (http://broadinstitute.github.io/picard/). Replicate bam files were merged and indexed using samtools. To obtain bedGraph files containing per-base CpG methylation metrics, MethylDackel (v0.3.0) was applied [extract --mergeContext --minDepth 2] (https://github.com/dpryan79/MethylDackel). Mitochondrial regions were subsequently removed as well as blacklisted regions for mm10 using bedtools [intersect -v] (v2.29.2) (ENCODE Project Consortium, 2012; Quinlan & Hall, 2010). To obtain bigwig files for genome browser visualization, bedGraph files were converted using the UCSC software bedGraphToBigWig (v4) (Kent *et al*, 2010).

#### Quantification of WGBS data at promoters

In order to quantify the percentage of CpG-methylation in promoter regions, bedGraph files were loaded into Rstudio using the bsseq package (v1.32.0) (Hansen *et al*, 2012). Subsequently, CpG-methylation was calculated in promoters of 3’ and promoter overlapping TUs and controls analogous to the CUT&Tag analysis (-500 bp to +200 bp around the TSSs) with options [getMeth(what = “perRegion”)]. Promoters without a called CpG were excluded from the analysis (<0.01% of all promoters). The resulting percentages were then visualized using the tidyverse package (v1.3.2) (Wickham *et al*, 2019) as a split violin plot.

### Experimental Methods

#### Cell lines

All experiments were performed in the female TXΔXic_B6_ line (clone A1) which is a F1 hybrid ESC line derived from a cross between the 57BL/6 (B6) and CAST/EiJ (Cast) mouse strains that carries a 773 kb deletion around the *Xist* locus on the B6 allele (chrX:103,182,701-103,955,531, mm10) and an rtTA insertion in the *Rosa26* locus (Pacini *et al*, 2021). The cell line was chosen, since it promotes Dox-inducible expression, but the Dox-inducible promoter present at the *Xist* gene in the parental line (TX1072) has been removed as part of the 773kb deletion. Moreover, the TXΔXic_B6_ line had been used to generate a large data set in a previous study that was used in the genome-wide analyses.

#### mESC culture and differentiation

mESCs were grown on 0.1% gelatin-coated flasks in serum-containing medium supplemented with 2i and LIF (2iL) (DMEM (Sigma), 15% ESC-grade FBS (Gibco), 0.1 mM β-mercaptoethanol, 1000 U/ml leukemia inhibitory factor (LIF, Millipore), 3 μM Gsk3 inhibitor CT-99021, 1 μM MEK inhibitor PD0325901, Axon). Differentiation was induced by 2iL withdrawal in DMEM supplemented with 10% FBS and 0.1mM β-mercaptoethanol on fibronectin-coated (10 μg/ml) tissue culture plates.

#### Molecular cloning

The lentiviral vector carrying the landing pad (SP419) and the donor plasmid with the antisense locus were generated by standard molecular cloning techniques (details are given in Suppl. Table 5). The annotated plasmid sequences are provided as Supplemental file 1. Starting plasmids were kind gifts from Hana El-Samad (MTK0_057, MTK0_017) (Fonseca *et al*, 2019) and from Luca Giorgetti (#235_237_pUC19_Chr1_Ho_CuO_Cre_HyTK). Correct identity of all plasmids was confirmed using Sanger sequencing. A list of primers used for cloning is also provided in Suppl. Table 5.

#### Generation of the TxSynAS line carrying a synthetic antisense construct

To generate the landing pad cell line TX_LVLP (SC55), TXΔXic_B6_ mESCs were transduced sequentially with a lentiviral vector encoding an ERT2-Gal4 construct for inducible expression (SP265), and the landing pad construct (SP419) (Fonseca *et al*, 2019). The ERT2-Gal4 is not used in this study.

##### Lentiviral transduction of the landing pad

For lentiviral transduction, 1*10^6^ HEK293T cells were seeded into one well of a 6-well plate and transfected the following day with the lentiviral packaging vectors: 1.2 μg pLP1, 0.6 μg pLP2 and 0.4 μg pVSVG (Thermo Fisher Scientific), together with 2 μg of the desired construct using Lipofectamine 2000 (Thermo Fisher Scientific). HEK293T supernatant containing the viral particles was harvested after 48 h. 0.2*10^6^ mESCs were seeded in a well of a 12-well plate in 2iL and transduced the next day with 1ml of 5x concentrated (lenti-X, Clontech) and filtered viral supernatant with 8 ng/μl polybrene (Sigma Aldrich). Puromycin (1 ng/μl, Sigma Aldrich) or hygromycin (200µg/ml, VWR) selection was started two days after transduction and kept until the selection control was dead but for at least 2 passages. To expand single clones, the cells were seeded at low densities in gelatin-coated 10 cm plates in 2iL medium and cultured until single colonies were visible (10 days). Then individual clones were picked and expanded. For the TX_LVLP line we selected clones that displayed high and homogenous expression upon BxB1-mediated integration of a GFP transcription unit (clones B1, A2).

##### BxB1-mediated integration

The TxSynAS line was then generated from the TX_LVLP line (clone B1) by insertion of the synthetic antisense locus into the landing pad through BxB1-mediated integration. To this end, the antisense plasmid (SP505) and a BxB1-encoding plasmid (SP225, Addgene #51271) were transfected with Lipofectamine 3000 (Thermo Fisher Scientific) in a 3-to-1 target-to-integrase ratio. 48h post transfection, blasticidin selection (5 ng/μl, Roth) was started, and single clones were expanded while selection was maintained. Clone C6 was chosen for all further experiments since it displayed high and homogenous GFP expression in the absence of doxycycline.

#### Memory experiment

For experiments, where reporter fluorescence was quantified by flow cytometry, all media were supplemented with 1µM Shield1 ligand (Takara, #632189) to stabilize both reporters, because the signals were too weak in the destabilized state without Shield1. 0.3*10^6^ cells, grown in 2iL, were seeded in a well of a 6-well plate in media containing 0µg/ml or 2µg/ml doxycycline, either in undifferentiated state (2iL, gelatine-coated culture dish) or with inducing differentiation (-2iL, fibronectin-coated culture dish). After two days, cells were harvested and seeded at a density of 7500 cells per well in a 96-well plate into media containing different Dox concentrations (0, 0.2, 0.4 or 2µg/ml), and assayed by flow cytometry 0h, 8h, 1d, 2d, 3d, and 4d later. To estimate background from autofluorescence a negative control (landing pad cell line w/o antisense construct integration) was measured in parallel in 2iL or -2iL.

#### Flow cytometry

Cells were analyzed using the BD FACSCelesta flow cytometer (Beckton Dickinson, IC-Nr.: 68186, Serial-Nr.:R66034500035) with 2-Blue6-Violet4-561YG laser configuration, and equipped with a BD High Throughput Sampler. The sideward and forward scatter areas were used for live cell gating. The height and width of the forward scatter were used for singlet/doublet differentiation. At least 10,000 events were recorded per replicate, initial condition, and doxycycline concentration. Fcs files were gated using RStudio with the *flowCore* (v1.52.1) and *openCyto* packages (v1.24.0) (Finak et al., 2014; Hahne et al., 2009). The mean fluorescence intensity (MFI) of the singlets was calculated. Background correction was performed by subtracting the MFI of the TX_LVLP line in the respective media.

#### WGBS

TXΔXic_B6_ were differentiated for 0, 2 or 4 days and genomic DNA was extracted using the DNeasy Blood and Tissue Kit (Qiagen). Bisulfite converted DNA libraries were prepared using the Accel-NGS Methyl-Seq DNA library kit (SWIFT BIOSCIENCES). In brief, 200ng (in 50µl lowTE) of purified DNA were fragmented to ∼350bp using the Covaris S2 system (10% duty cycle, intensity 5 for 2x 45 sec in Covaris AFA tubes) followed by a concentration step with Zymo columns (DNA Clean & Concentrator). 20µl of DNA were used for bisulfite conversion over night with the Zymo EZ-DNA methylation Gold Kit.

Bisulfite converted DNA was fully denatured by 2 min incubation at 95°C and immediately transferred to ice. To anneal a truncated adapter, 15µl of denatured DNA was mixed with 25µl of Adaptase reaction mix (SWIFT) and incubated for 15min at 37°C, 2min at 95°C, and then cooled to 4°C. Extension and second strand synthesis by using a primer complementary to the truncated adapter was performed mixing the Adaptase reaction mix with 44µl of extension reaction mix (SWIFT) and incubation at 98°C for 1min, 62°C for 2min, 65°C for 5min and cooling to 4°C. The samples were cleaned with 1.2 volumes of Ampure XP beads, 80% ethanol and eluted in 15µl. Ligation of the second (truncated) adapter was performed at 25°C for 15min followed by an additional bead clean up with 1 volume of AmpureXP beads and 80% ethanol. Samples were individually indexed using unique dual indexed primer sets with 6 cycles of a slightly modified PCR program (30s @ 98°C, 6 cycles of 15s @ 98°C, 30s @ 60°C, 60s @ 68°C, followed by a final 5 minutes incubation step at 68°C. Libraries were finally cleaned up with 1 volume of Ampure XP beads. Quality was assessed using Agilent’s Bioanalyzer and concentration was determined by qPCR. Libraries were pooled equimolarly and sequenced on a NovaSeq 6000 S2 flowcell (Illumina) in paired end 150 mode to yield ∼120-278mio fragments.

## Acknowledgements

We would like to thank the FACS facility, the sequencing facility and the IT service at the MPI for molecular Genetics for maintenance and support of a computing cluster. We thank Luca Giorgetti and Hana El-Samad for sharing plasmids. This work was supported by the Max-Planck Research Group Leader program, E:bio Module III—Xnet grant (BMBF 031L0072) and ERC Starting Grant CisTune (948771) to E.G.S. V.M. and T.S. were supported by the DFG (GRK1772, IRTG 2403). G.N. was supported by the European Union’s Horizon 2020 Research and Innovation Program (Marie Skłodowska-Curie ITN PEP-NET).

## Data and code availability

Code used in this study is available at https://github.com/EddaSchulz/Antisense_paper. DNA methylation data generated is available on GEO under accession number GSE253792.

## Materials

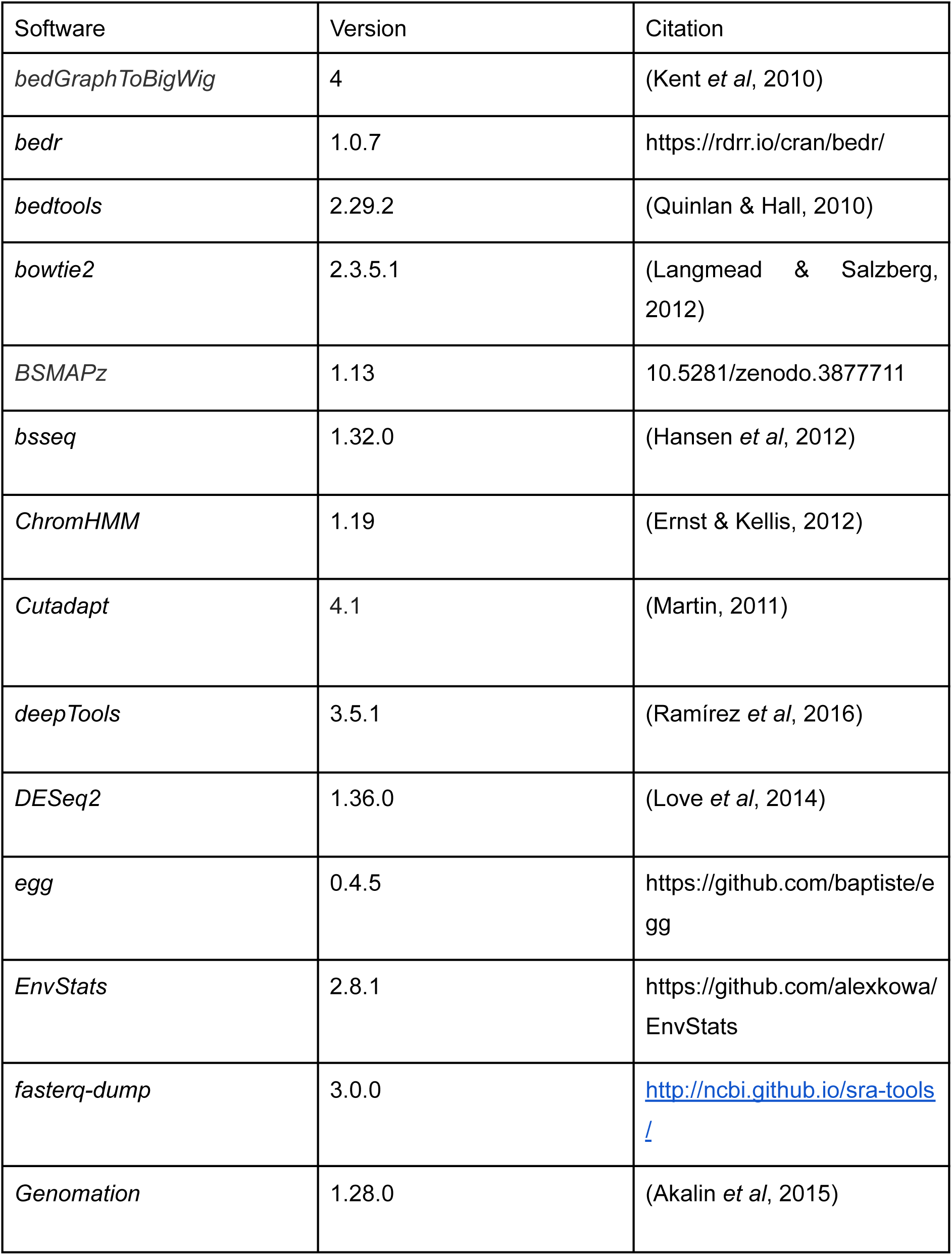
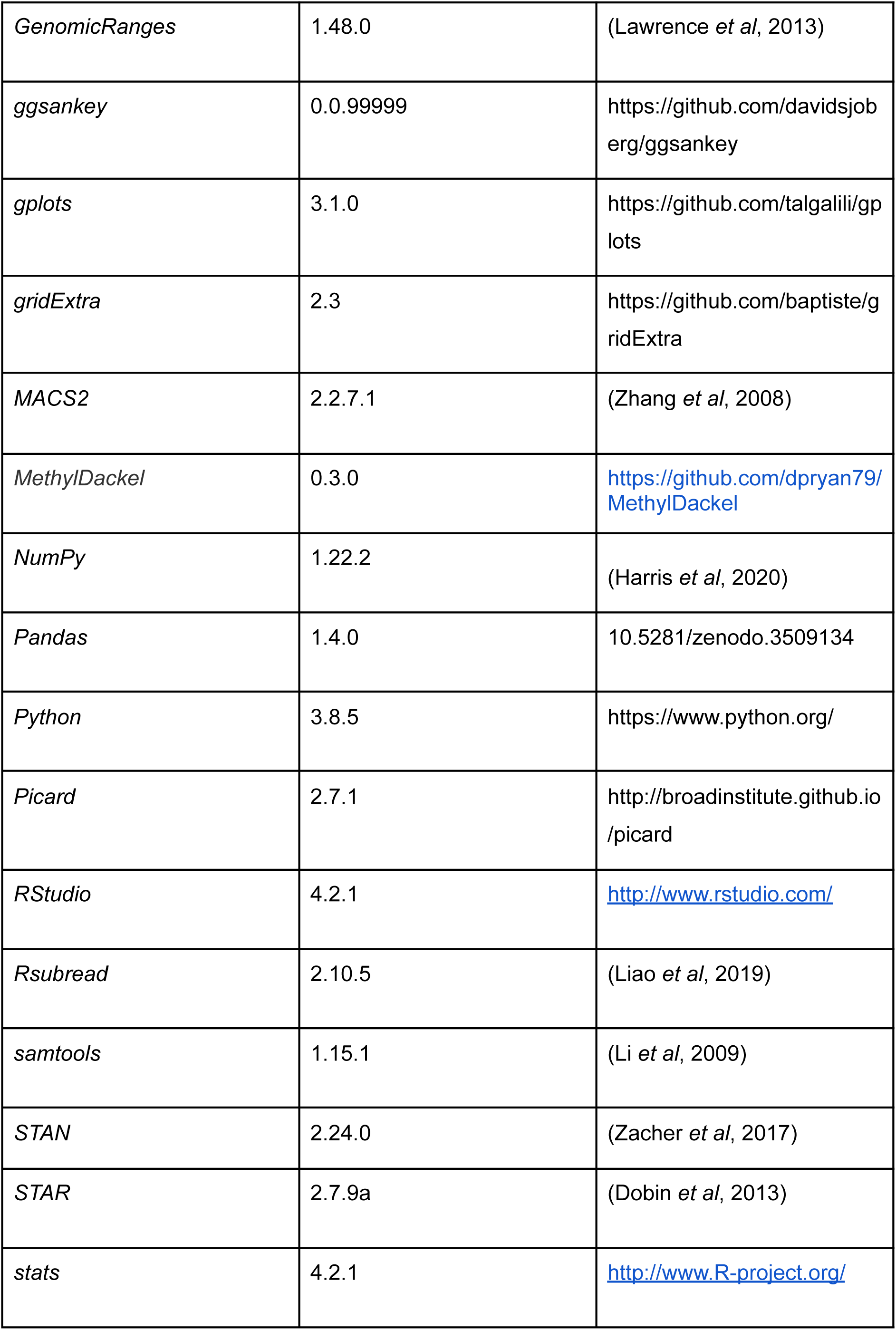
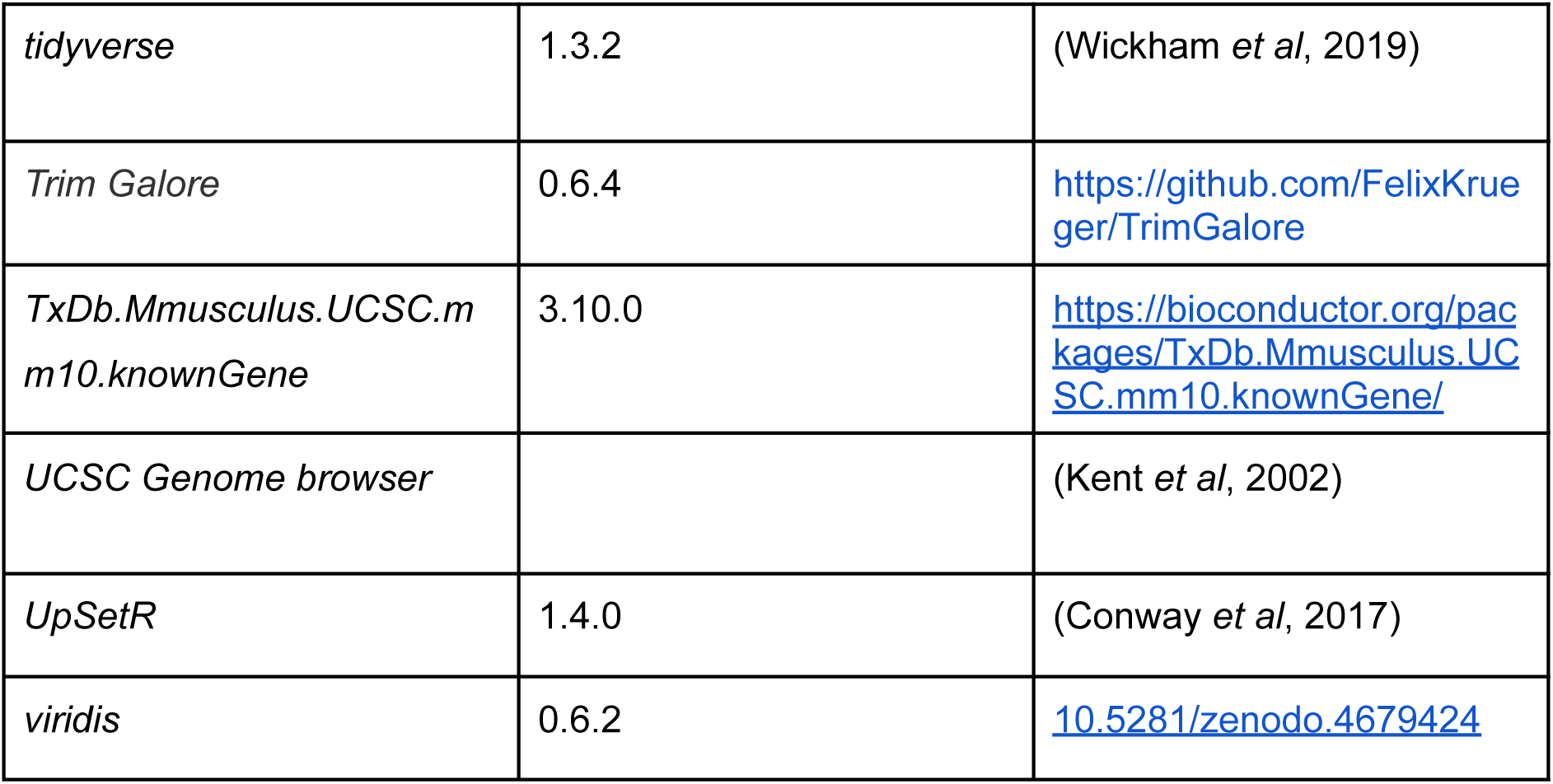

## Supplementary Figures

**Suppl. Fig. 1.**
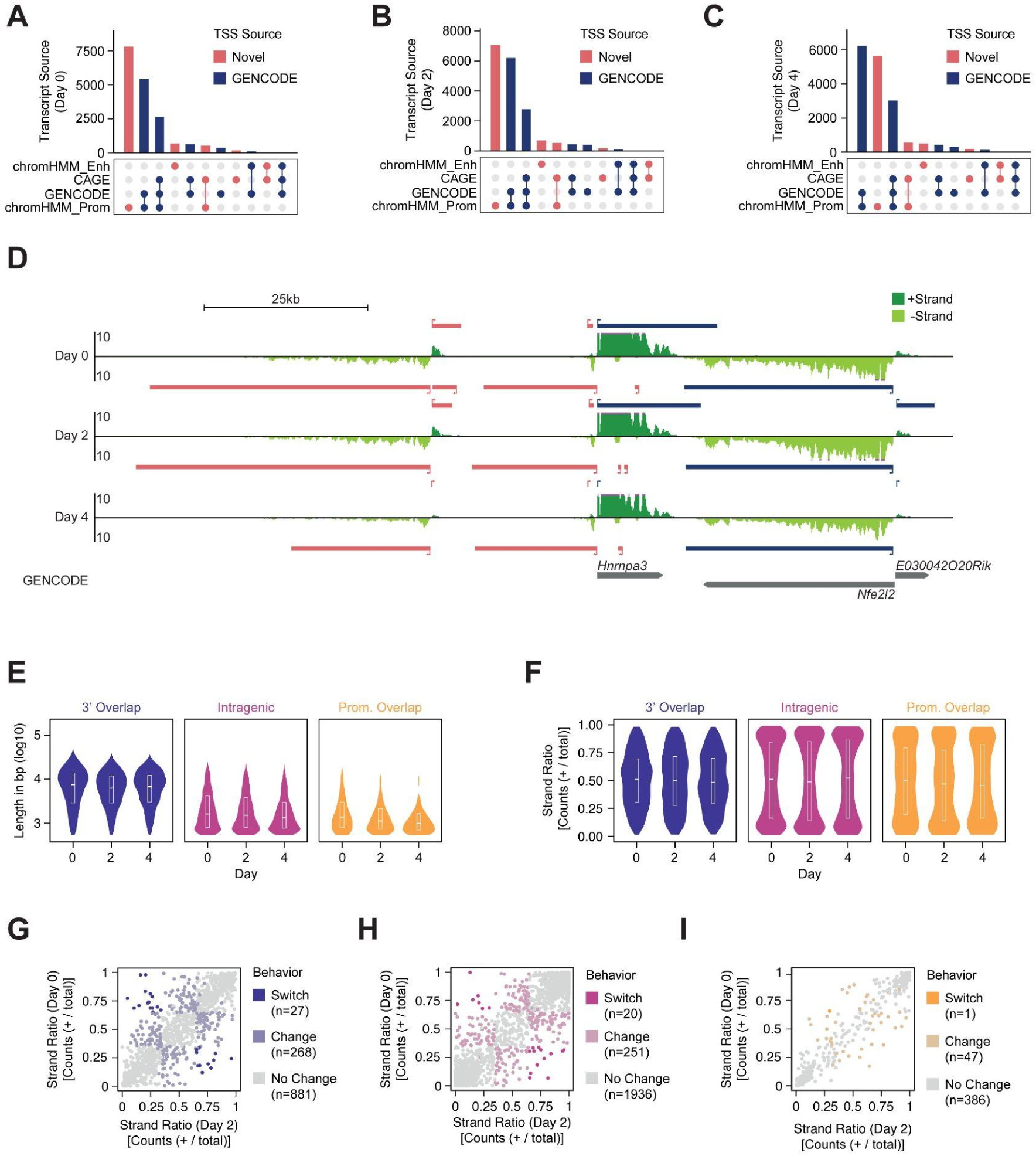
Genome-wide annotation of antisense transcription. *Related to* figure 3. A-C) UpSetR plots showing the sources for the TSSs of annotated TUs for day 0 (A), 2 (B) and 4 (C) of differentiation. TSSs with GENCODE annotations are shown in blue. Novel TSS are shown in red. D) Genome browser screenshot of an example locus with nascent transcriptome assembly. TUs on the plus strand are shown above and TUs on the minus strand below the tracks for each timepoint. Novel TUs are colored in red, while GENCODE transcripts are shown in blue. TT-seq signal that extends beyond the scale is marked with purple on the edge of the tracks. E) Length of identified overlaps separated by type at days 0, 2 and 4 of differentiation. F) Strand ratio within the different assemblies separated by day and overlap type. The ratio was calculated as the fraction of counts mapping to the plus strand. (G-I) Scatter plot comparing the strand ratio of 3’ overlaps (G), intragenic overlaps (H) and promoter overlaps (I) at day 0 and day 2. An overlapping region was categorized as biased, if transcription was significantly different between the plus and minus strands (FDR<=0.1, Student’s T-test with Benjamini-Hochberg correction) and the strand ratio was >= 0.65 (plus bias) or <= 0.35 (minus bias). An overlap was annotated as ‘switch’, if it changed from one bias to the other and as ‘change’ if it changed from no bias to bias (or vice-versa).

**Suppl. Fig. 2.**
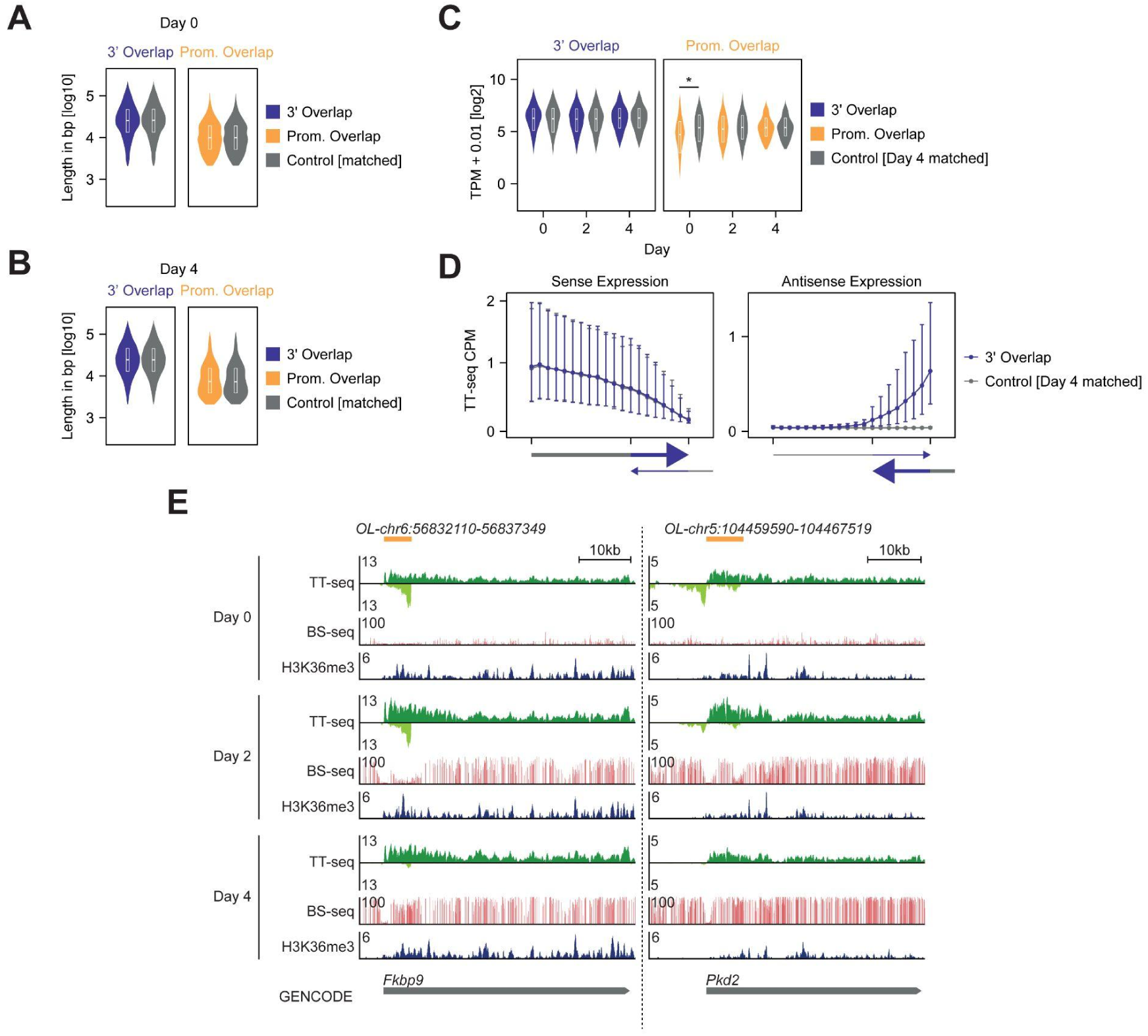
Systematic comparison of antisense loci with overlap-free genes. *Related to* figure 4. A) Length comparison between 3’/promoter-overlapping TUs and matched controls at day 0 (A) or 4 (B). C) Expression measured by TT-seq of 3’ and promoter-overlapping transcribed regions, annotated based on nascent transcription at day 4 of differentiation in comparison to matched controls. Significance was assessed using a ranked Wilcoxon sum test (p<=0.01). D) Binned line plot showing sense or antisense expression of TT-seq data in 3’ overlap transcribed regions and controls at day 4. Expression was quantified in 12 “free” bins and 8 “overlapping” bins. The big dots depict the median of all transcribed regions, while the upper and lower whiskers depict 3rd and 1st quartiles respectively. Significance was assessed using a ranked Wilcoxon sum test (p<=0.01). E) Genome browser screenshot showing TT-seq, BS-seq and H3K36me3 CUT&Tag data for two promoter overlaps at which transcription correlates with DNA-methylation. TT-seq reads from the plus-strand are colored in dark green, while reads from the minus-strand are shown in light green. The location of the overlaps is shown above the tracks.

**Suppl. Fig. 3.**
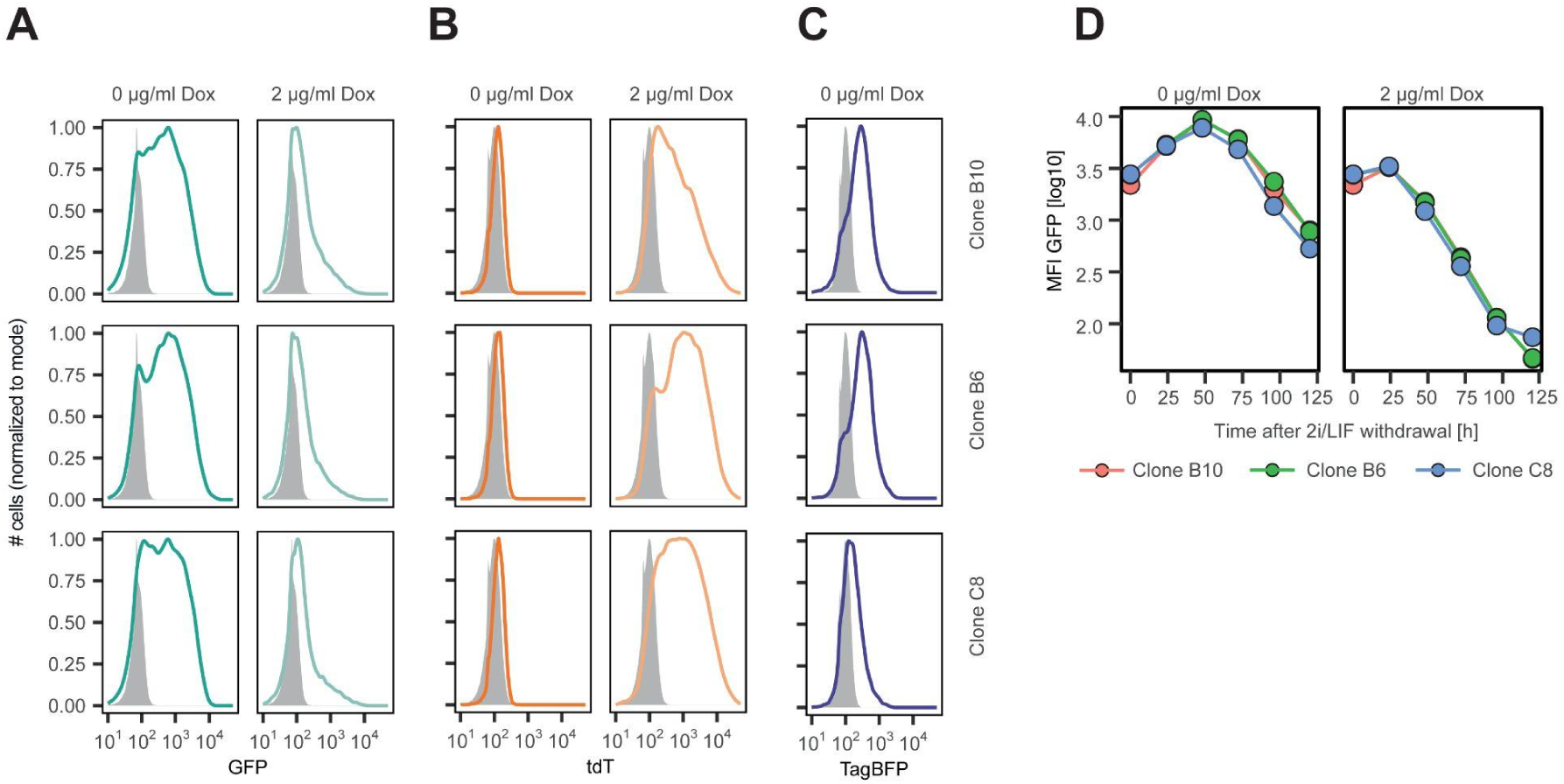
GFP expression in TxSynAS cells is down-regulated upon differentiation independent of antisense transcription. *Related to* figure 5 A-C) GFP (A), tdTomato (B) and TagBFP (C) expression in three additional clones of the TxSynAS line, where the synthetic locus was integrated in two different genomic positions, cultured for 2 days in the presence (light colors) or absence (dark colors) of Dox. TagBFP is expressed only upon integration into the landing pad. D) GFP expression upon differentiation by 2i/LIF withdrawal in single clones of the TxSynAS line with different genomic antisense integration sites in the presence of 0µg/ml (left) and 2µg/ml (right) doxycycline.

## Supplementary Tables

**Suppl. Table 1**

Simulations of antisense transcription: parameter ranges tested and statistics of FST analysis (related to Fig. 1-2)

**Suppl. Table 2**

Genome-wide annotation of antisense transcription: De novo annotation of transcribed regions; detected antisense loci; transcriptional activity of all transcribed regions, and of antisense loci; composite antisense loci; switching antisense loci (related to Fig. 3)

**Suppl. Table 3**

Characterization of antisense loci: Control regions used for comparison; statistical analyses comparing overlapping, and non-overlapping control regions (related to Fig. 4)

**Suppl. Table 4**

Statistical analysis of experimentally measured memory in mESC conditions, based on the ODE model analysis (related to Fig. 5)

**Suppl. Table 5**

Molecular Cloning: cloning strategies plasmids and primers

## Supplementary Files

### Suppl. File 1

Annotated sequences of plasmids used in the study (SP505, SP419) in Genebank format.

